# Paralog diversification masks conserved diel regulatory programs during cold acclimation in *Brassica rapa*

**DOI:** 10.64898/2026.07.24.740384

**Authors:** Angela Ricono, Zachary A. Myers, Danielle Schoenecker, Ananda Menon, Dennis Such, Adelaide Hazen, Ava Wise, Tomáš Brůna, Jerry Jenkins, Christopher Plott, Jenell Webber, LoriBeth Boston, Shengqiang Shu, Yinjie Qiu, Kerrie Barry, Chiemeka K. Nwakama, Jane Grimwood, Jeremy Schmutz, John T. Lovell, Kathleen Greenham

## Abstract

Plant stress responses occur within daily cycles of physiology, metabolism, and growth, making timing a critical dimension of acclimation. In Arabidopsis, circadian and diel regulation influence responses to abiotic stress, including cold, but how this temporal regulation is conserved, diversified, or expanded in crop genomes remains unclear. This question is especially challenging in *Brassica rapa*, which underwent a genome triplication after diverging from Arabidopsis, resulting in multiple retained paralogs that can be grouped by Arabidopsis orthology and ancient homeologous relationships. Here, we generated a *B. rapa* pangenome spanning six morphotypes and used it to profile diel (24 h) cold acclimation responses across diverse accessions differing in freeze tolerances. Cold altered peak expression time for thousands of genes, which we grouped into distinct phase-change groups. Circadian leaf movement assays revealed accession-specific differences in clock period and temperature compensation under cold, suggesting that altered clock behavior may contribute in part to the diel transcriptome retiming. At the individual gene level, inferred gene regulatory networks (GRNs) were highly accession-specific and lost shared connectivity under cold stress. However, grouping these paralogs by their Arabidopsis orthologs revealed a highly conserved regulatory architecture that was otherwise masked by paralog diversification. Integrating these networks with functional pathways identified key candidate regulators of retimed processes, including modules linked to nighttime phosphorylation and daytime photosynthesis. Finally, analyzing conserved noncoding sequences across the pangenome prioritized specific regulatory targets within cold-retimed groups. Together, these results demonstrate that cold acclimation in B. rapa is shaped by a combination of diel retiming, paralog-specific regulation, and deeply conserved programs.

## Introduction

Extreme temperature events, including cold and freezing episodes, significantly hinder crop productivity and are expected to intensify with climate instability (Soualiou et al. 2022; Fanzo et al. 2025; Feng et al. 2025). Cold tolerance reflects the combined effects of pre-existing traits and inducible acclimation responses such as variation in membrane lipid composition, accumulation of protective metabolites, and altered gene expression programs (Thomashow 1999; Cook et al. 2004; Lissarre et al. 2010; Degenkolbe et al. 2012; Shomo et al. 2024). Improving crop resilience to low-temperature stress therefore requires a deeper understanding of the genetic and regulatory mechanisms that enable plants to acclimate to cold and survive subsequent freezing. In crop species with extensive genomic and morphological diversity, natural variation is an important resource for comparative studies aimed at identifying these mechanisms (Liu et al. 2014; Golicz et al. 2016; Soltis and Soltis 2016; Gabur et al. 2019; Schiessl et al. 2019; Cai et al. 2021; Qi et al. 2021; Wang et al. 2025b).

Structural variation, including presence absence variation, copy-number variation, differential fractionation, and relative rearrangement, contributes substantially to phenotypic diversity in crops (Würschum et al. 2015; Schiessl et al. 2019; Wang et al. 2025b). Much of this diversity is poorly represented when using a single reference genome but can be captured through pangenomic approaches. Pangenomes incorporate genomes from multiple individuals within a species, representing the entire breadth of genes that are present among genetically similar populations, and allow us to capture both core (conserved among genomes) and accessory (variable retention among genomes) genes (Golicz et al. 2016, 2020; Cai et al. 2021; Hufford et al. 2021; Li et al. 2023; Zhao et al. 2025).

Across crop species, accessory genes and copy-number variable regions are frequently associated with functions related to defense, signaling, flowering, fruit quality/ripening, specialized metabolism, and abiotic stresses including freezing tolerance (Hofberger et al. 2013; Würschum et al. 2015; Golicz et al. 2016, 2020; Panchy et al. 2016; Montenegro et al. 2017; Hurgobin et al. 2018; Alonge et al. 2020; Song et al. 2020; Cai et al. 2021). For example, glucosinolate biosynthetic genes, which encode sulfur-rich specialized metabolites with well established roles in biotic and abiotic stress tolerance (Hofberger et al. 2013; Han et al. 2023), show extensive copy-number variation across Brassicaceae (Zang et al. 2009; Hofberger et al. 2013; Golicz et al. 2016; Abrahams et al. 2020) and contribute to differences in glucosinolate content (Zhang et al. 2018; Wei et al. 2019), defense responses (Han et al. 2023), and sulfur allocation (Sugiyama et al. 2021). Likewise, variation in *FLOWERING LOCUS C* (*FLC*) is associated with differences in vernalization requirement and flowering time among Brassica morphotypes (Zhao et al. 2010; Golicz et al. 2016; Calderwood et al. 2021; Akter et al. 2023). These examples demonstrate that genomic variation can shape adaptive traits relevant to environmental resilience (Golicz et al. 2016).

*Brassica rapa* (*B. rapa*) is an especially useful system to examine how genomic diversity contributes to abiotic stress adaptation. Its close evolutionary relationship with economically important crops such as Canola, as well as the model plant Arabidopsis, further positions it as an attractive model for identifying conserved and accession-specific mechanisms of cold acclimation and freezing tolerance. Following divergence from Arabidopsis approximately 20 MYA (Hohmann et al. 2015), the *B. rapa* lineage underwent a whole-genome triplication followed by extensive genome fractionation, a process involving biased loss/retention of duplicated genes, leading to substantial paralog diversity and structural variation (Wang et al. 2011; Cai et al. 2021). *B. rapa* encompasses numerous specialty crops (*e.g.*, chinese cabbage, pak choi, oilseed, turnip and leafy vegetable varieties) that vary dramatically in their leafy-head architecture, storage-root formation and oilseed forms (Cheng et al. 2014; Bird et al. 2017; Lu et al. 2025). The diversity of form in *B. rapa* is also due to the unique domestication histories of morphologically distinct cultivars (herein referred to as ‘morphotypes’) (McAlvay et al. 2021; Cai et al. 2026). Historical and population genomic analyses support multiple domestication trajectories, with a model describing an early Eurasian-East Asia origin generally accepted (Guo et al. 2014; Bird et al. 2017; Qi et al. 2017). *B. rapa* crops were domesticated across temperate, arid, and semi-arid regions, suggesting that artificial and environmental selection may have acted on distinct abiotic stress responses across lineages. (Mäkelä et al. 2011; Qi et al. 2017).

A common environmental stress for temperate crops including Brassicas is exposure to low, non-freezing temperatures, often followed by subsequent freezing events, which are mitigated through a process known as cold acclimation (Ding et al. 2019). Cold acclimation is a complex process that, following exposure to non-lethal low temperatures, increases tolerance to subsequent freezing events (Thomashow 1999; Ding et al. 2019). The acclimation process broadly involves the coordination of hormonal and transcriptional cascades that remodel membrane lipids to maintain structural fluidity (Shomo et al. 2024), increase antioxidant pools (e.g., glutathione, anthocyanins) (Li and Ahammed 2023; Qian et al. 2024), and increase intracellular osmolyte content (e.g., proline, sugars, carbohydrates) (Thomashow 1999), to regulate water homeostasis (Lissarre et al. 2010; Ding et al. 2019; Qian et al. 2024). After exposure to sub-zero temperatures, ice forms outside the cell, reducing extracellular water potential and effectively dehydrating the cell as water flows out due to osmosis (Thomashow 1999; Xin and Browse 2000; Shomo et al. 2024).

Although the earliest mechanisms of cold perception remain incompletely resolved, low temperature rapidly rigidifies membranes and induces cytosolic calcium influx (Thomashow 1999), triggering signaling cascades that activate cold responsive transcriptional regulators including MYB, bZIP, NAC, and WRKY transcription factor (TF) families (Wang et al. 2014; Chen et al. 2015; Liu et al. 2021; Abdullah et al. 2022). Central to this response is the ICE-CBF-COR pathway in which the bHLH factor Inducer of CBF Expression 1 (ICE1) and CALMODULIN-BINDING TRANSCRIPTION ACTIVATOR (CAMTA) promote induction of C-repeat binding factors/Dehydration-responsive element binding AP2/ERF family (CBFs/DREBs), alongside additional cold-responsive TF families (Ding et al. 2019; Abdullah et al. 2022). Together, these TFs activate large sets of cold-responsive (*COR*) genes (Shi et al. 2017). However, the contribution of individual pathway components varies by species, accession, and paralog (Song et al. 2020; Deng et al. 2024). In Arabidopsis, variation at *CBF* loci contributes to different freezing tolerances among accessions (Gehan et al. 2015). Comparative transcriptomic studies in winter rapeseed have shown that key cold-responsive TFs, including *ICE1* paralogs (Wu et al. 2024) and numerous *WRKY* TFs (Kayum et al. 2015), are differentially induced among freezing-tolerant versus freezing-sensitive *B. rapa* lines (Wu et al. 2022). Transgenic studies in wheat show *TaCBF14* and *TaCBF15* homologs improve freezing tolerance by upregulating expression of downstream *COR* genes (Soltész et al. 2013), while other *CBF/DREB* homologs contribute to cold tolerance in rice (Ito et al. 2006; Zhang et al. 2009; Moon et al. 2019) and canola (Savitch et al. 2005) by upregulating expression of additional *COR* or photosynthesis genes (Savitch et al. 2005; Zhang et al. 2009; Dahal et al. 2012). These findings support that paralog diversification is an important source of variation in cold acclimation, particularly in polyploid crops.

Because acclimation responses can be metabolically costly, the magnitude and timing of acclimation are strongly time-of-day dependent and are regulated in part by the circadian clock, particularly through clock control of the CBF regulon (Dong et al. 2011; Maibam et al. 2013). The circadian clock is an endogenous ∼24h oscillator composed of interlocked transcriptional-translational feedback loops which, in plants, are primarily entrained by light and temperature (Harmer 2025). Circadian regulation of the transcriptome enables temporal allocation of resources and responsiveness to the times of day when they are most beneficial, a phenomenon referred to as circadian gating (Graham et al. 2023). In Arabidopsis, the clock gates cold responsiveness in part through morning expressed regulators, CIRCADIAN CLOCK ASSOCIATED 1 (CCA1) and LATE ENLONGATED HYPOCOTYL (LHY), which restrict the activity of the CBFs and other downstream CORs (*i.e.* COR15A; COR47; COR78) (Dong et al. 2011) to specific times of day. Other cold-responsive TFs are under similar time-of-day regulation (*i.e.* RAV1; ZAT12) (Fowler et al. 2005), indicating that cold response pathways are integrated within the circadian network. In *B. rapa,* natural variation in the clock-associated gene *GIGANTEA* (*GI*), a key regulator of the clock and photoperiodic signalling (Nohales et al. 2019), is associated with differences in circadian period in *B. rapa*, and these differences correlate with variation in cold and freezing tolerance (Xie et al. 2015). In Chinese cabbage, alleles of *EARLY FLOWERING 3* (*ELF3*), a well known clock component best described for integrating light and temperature signals into the clock (Ezer et al. 2017), alter thermal responsiveness of circadian rhythms (Wang et al. 2025a). Moreover, clock genes have been preferentially retained in multiple copies in *B. rapa*, and retained paralogs often exhibit divergent temporal expression patterns (Greenham et al. 2020).

These observations raise the possibility that time-of-day regulation and paralog diversification jointly shape cold acclimation responses. A major unresolved question is whether cold tolerance in *B. rapa* is driven by conserved regulation of shared pathways, morphotype-specific domestication histories, or accession-specific use of distinct paralogs within broadly conserved stress-response networks. Here, we combined time-of-day-resolved freezing assays with a cold acclimation RNA-seq time course across 13 *B. rapa* accessions representing six distinct morphotypes. We used these data to determine how cold acclimation reshapes diel transcriptional programs, and whether cold-responsive pathways are conserved at the paralog or ortholog level. By integrating temporal expression analysis with gene regulatory network inference, we identified both conserved and accession-specific regulatory features associated with cold acclimation. Our results show that *B. rapa* accessions undergo extensive transcriptional reprogramming under cold, but that freezing tolerance is not explained by the number of cold-responsive genes or by morphotype. Instead, accessions differ in the timing of pathway activation and in the specific paralogs recruited during acclimation. An examination of conserved noncoding sequences reveals insight into the altered regulatory landscape across *B. rapa* that has shaped these differential responses. These findings demonstrate that temporal regulation and genome evolution intersect to shape cold acclimation in a mesopolyploid crop and highlight the importance of incorporating time-of-day and pangenome approaches into strategies for improving freezing tolerance.

## Methods and Materials

### Plant growth and experimental design

Seeds for 14 *B. rapa* lines (Supplemental Table1, Accession Information tab) were germinated on wet whatman paper for three days before sowing in 3.5” by 3.5” by 3.5” (height; width; depth) rectangular pots filled with moistened Sungrow Professional Growing Mix soil. Plants were grown in 14h/10h light/dark photoperiods with 200-250 m^2^s^-1^ light intensity, 21°C/16°C temperature cycles, and 60-70% relative humidity. All plants were fertilized with 30mL of 0.5X Hoaglands liquid fertilizer 14 days after planting. Ten replicates per accession were cold acclimated after three weeks of growth in control conditions, with an additional five replicates per line for electrolyte leakage assays. Cold acclimation conditions for the first day consisted of a six hour ramp down from 21°C day to a 4°C night. Temperatures increased to 10°C at ZT0 (Day 2; ZT: Zeightgeber or hours after lights on) and were ramped back down to 4°C for eight hours at ZT13, followed by a freeze ‘snap’ at -4°C for two hours. Temperatures were brought back up to -2°C at ZT0 for two hours (Day 3) then slowly warmed to 4°C and maintained at 4°C for the remainder of the day. Nighttime temperatures were slowly ramped up from 4°C to 16°C (end of Day 3) over the course of 10 hours. Temperatures were then slowly ramped from 16°C (nighttime control conditions) to 21°C (daytime control conditions) over the course of eight hours (Day 4). These conditions were maintained for the remainder of the experiment.

### Physiology assays following freezing stress

To track alterations to physiology throughout the experiment we measured several aspects of plant physiology before (Day 1), during (‘Acclimation’ or Day 2; ‘Freeze’ or Day 3; and ‘Post-Freeze’ or Day 4) and after (Recovery on Days 5, 9 and 10) cold acclimation. These metrics include light adapted photosystem II (PSII) efficiency (Fv’/Fm’), leaf damage, electrolyte leakage, and above ground (fresh) biomass. PSII efficiency was measured on ten biological replicates using a handheld fluorometer (‘FluorPen’) at various timepoints throughout the experiment. Leaf tissue from five biological replicates were collected for electrolyte leakage at ZT0 the day following the freeze (Day 4). To avoid cell rupture that might occur by attempting to submerge a whole leaf in a small test tube, we cut one half-inch by half-inch section on the left and right side of the midvein for each leaf using a sharp razor blade. Sections were fully submerged in 4mL of DI water in glass test tubes and shaken for three hours before measuring the initial conductivity. Leaf sections without DI water were then frozen at -80℃ overnight to rupture the remaining intact cells before measuring the final conductivity as described above. Three conductance readings per leaf section were taken and then averaged for each section. The average leakage was calculated as between the two leaf sections and used for statistical analyses and plotting. To assess both freezing damage and recovery capacity, we calculated the percent change in post-freeze and recovery measurements relative to the pre-acclimation levels. Leaf damage was assessed on ten biological replicates by giving each plant a ‘damage score’ which is a whole number rating that describes the percentage of damage observed, where ‘10’ is no damage to any leaves, ‘1’ is a fully dead plant, and ‘5’ is approximately 50% of the plant exhibiting damage. Above ground biomass was collected on five biological replicates on the tenth and last day of the experiment. These data (R_input/BrapaPhys.RData) are provided and the code to analyze and visualize them (<u>BrapaColdAcclimation.R</u>) are provided on our Github (https://github.com/greenhamlab/B.rapa-diel-cold-regulation/).

### Nucleic Acid extractions

High molecular weight DNA was extracted from young tissue using the protocol of Doyle and Doyle (Doyle and Doyle. 1987) with minor modifications. Flash-frozen young leaves were ground to a fine powder in a frozen mortar with liquid nitrogen followed by very gentle extraction in 2% CTAB buffer (that included proteinase K, PVP-40 and beta-mercaptoethanol) for 30min to 1h at 50 °C. After centrifugation, the supernatant was gently extracted twice with 24:1 Chloroform : Isoamyl alcohol. The upper phase was transferred to a new tube and 1/10th volume 3 M Sodium acetate was added, gently mixed, and DNA precipitated with iso-propanol. DNA precipitate was collected by centrifugation, washed with 70% ethanol, air dried for 5-10 minutes and dissolved thoroughly in elution buffer at room temperature followed by RNAse treatment. DNA purity was measured with Nanodrop; DNA concentration was measured with Qubit HS kit (Invitrogen), and DNA size was validated by Femto Pulse System (Agilent).

### Genome assembly

The seven genome assemblies were constructed with identical pipelines and identified only minor differences in coverage and support (Supplemental Table2). Initial assemblies were constructed with PacBio HiFi and Dovetail OmniC Hi-C data and assembled using HiFiAsm+HIC (Cheng et al. 2021). The initial assembly was polished with RACON (Vaser et al. 2017). Contigs for both haplotypes were oriented, ordered, and joined with JUICER (Durand et al. 2016). Contigs containing significant telomeric sequence were considered properly oriented in the assembly. All initial assemblies were manually screened to identify misjoins using Hi-C data. Any misjoins present were corrected. Where necessary, adjacent alternative haplotypes were identified on the joined contig set and collapsed using the longest common substring between the two haplotypes. Homozygous SNPs and INDELs were identified with 2×150, 400bp insert Illumina reads mapped to each reference with GATK and corrected in the release sequence (McKenna et al. 2010). Chromosomes were numbered and oriented using version 1.0 Brassica rapa var. FPsc (Turnip mustard) obtained from Phytozome (https://phytozome-next.jgi.doe.gov). The resulting sequence was screened for retained vector and/or contaminants.

### Genome annotation

Transcript assemblies were built using PERTRAN (Lovell et al. 2018), which conducts genome-guided transcriptome short read assembly via GSNAP (Wu, 2010) and builds splice alignment graphs after alignment validation, realignment and correction. To obtain putative full length transcripts, PacBio Iso-Seq CCSs were corrected and collapsed by genome guided correction pipeline, which aligns CCS reads to genome with GMAP (Wu, 2010) with intron correction for small indels in splice junctions if any and clusters alignments when all introns are the same or 95% overlap for single exon. Subsequently transcript assemblies were constructed using PASA (Haas, 2003) from RNA-seq transcript assemblies. Loci were determined by transcript assembly alignments and/or EXONERATE alignments of proteins from Arabidopsis (Arabidopsis thaliana), soybean, poplar, cotton, rice, sorghum, peach, citrus, aquilegia, tomato, grape, *Eutrema salsugineum*, *Nymphaea colorata*, *Amborella trichopoda*, and Swiss-Prot proteomes to repeat-soft-masked assemblies using RepeatMasker (Smit, 2013-2015) with up to 2K BP extension on both ends unless extending into another locus on the same strand. Gene models were predicted by homology-based predictors, FGENESH+ (Salamov, 2000), FGENESH_EST (similar to FGENESH+, but using EST to compute splice site and intron input instead of protein/translated ORF), EXONERATE (Slater, 2005), PASA assembly ORFs (in-house homology constrained ORF finder), and AUGUSTUS (Stanke, 2006) trained by the high confidence PASA assembly ORFs and with intron hints from short read alignments. The best scored predictions for each locus are selected using multiple positive factors including EST and protein support, and one negative factor: overlap with repeats. The selected gene predictions were improved by PASA. Improvement includes adding UTRs, splicing correction, and adding alternative transcripts. PASA-improved gene model proteins were subject to protein homology analysis to above mentioned proteomes to obtain Cscore and protein coverage. Cscore is a protein BLASTP score ratio to MBH (mutual best hit) BLASTP score and protein coverage is the highest percentage of protein aligned to the best of homologs. PASA-improved transcripts were selected based on Cscore, protein coverage, EST coverage, and their CDS overlapping with repeats. The transcripts were selected if their Cscore is larger than or equal to 0.5 and protein coverage larger than or equal to 0.5, or it has EST coverage, but their CDS overlapping with repeats is less than 20%. For gene models whose CDS overlaps with repeats for more than 20%, their Cscore must be at least 0.9 and homology coverage at least 70% to be selected. The selected gene models were subject to Pfam analysis and gene models whose protein is more than 30% in Pfam TE domains were removed and weak gene models. The genome is hardmasked with its high confidence gene models produced above (transcriptome fully supported, high homology supported and complete gene models). The masked genome was BLASTXed and EXONERATEed against non-self high confidence and non-redundant peptide proteomes from 7 genomes (MIZ_19 of B. juncea, and O_302V, L58, PCGlu, A03, VT123, and WO_83 of B. raps) in Brassica pan-genome annotations to make EXONERATE gene predictions. The predicted gene models were scored with BLASTP using homology proteomes. These models were compared to ones from the first round and better homology supported ones (e.g., not contra-indicated by transcriptome evidence) were kept to improve gene models from the first round or added if not found therein. Incomplete gene models, low homology supported without fully transcriptome supported gene models and short single exon (< 300 BP CDS) without protein domain nor good expression gene models were manually filtered out.

### GENESPACE analyses

To identify syntenic relationships among the *B. rapa* lines we used the R-package GENESPACE (Lovell et al. 2022). We also included Arabidopsis in a separate GENESPACE run for comparative analyses and to identify paralogs within the morphotypes. Following this we employed an in-house script to assign Brassica genes that were assigned to multiple Arabidopsis orthologs to a single ortholog. We repeated this framework for Arabidopsis genes that may have been assigned to multiple Brassica genes. Finally, to account for tandem duplications which are known to be relatively predominant in polyploids, we outline a multi-step process to ensure that all syntenic tables reflect one-to-one relationships between Brassica annotations and Arabidopsis. Scripts to create the syntenic tables are available on our GitHub (https://github.com/greenhamlab/B.rapa-diel-cold-regulation).

### Plant growth and experimental design for cold acclimated RNA sequencing

Plants from 14 *B. rapa* accessions were grown for three weeks in in (3.25’ x 3.625’) pots with a soil mixture of two parts Metro-Mix PX1 + one part Pro-Mix amended with 0.5 mL of Osmocote 18-6-12 fertilizer (Scotts, Marysville, OH) under 24/14℃ thermal cycles and 14h /10h light/dark cycles. Plants were cold acclimated for three days starting with a drop in temperature at ZT14 (lights off) to 4℃, followed by an increase to daytime temperatures of 10℃ at ZT0. Whole tissue was collected for RNA sequencing starting at ZT17 on the third night of acclimation from four individual plants (biological replicates) per timepoint. Tissue was collected every four hours over a 24 hour period for a total of six timepoints. RNA extraction was performed at the Cornell Institute of Biotechnology using the Direct-zol-96-RNA extraction kit by Zymo (Irvine, CA) and sent to the Joint Genome Institute for sequencing as part of project CSP504418 (See below).

### RNA sequencing and processing of transcripts

RNA was submitted to the DOE Joint Genome Institute for library preparation and sequencing as part of a Community Science Program grant CSP504418. Paired-end RNA sequencing was performed on a NovaSeq. Data are available through the NCBI GEO database. Reference genomes are available on Phytozome (https://phytozome-next.jgi.doe.gov/). Raw RNA sequencing reads were filtered and trimmed using BBDuk (Bushnell 2014) and were mapped to every reference genome. Mappings with the highest score were used for downstream analyses. Libraries were aligned using HISAT2 v2.2.1 (Kim et al. 2015) and strand specific files were generated using deepTools v3.1 (Ramírez et al. 2016). Feature count tables were generated using featureCounts (Liao et al. 2014) from hits assigned to the reverse strand and were normalized to Counts Per Million (CPMs). Genes that did not have at least four samples greater than zero (log2 CPM) were removed. Missing replicates were imputed by averaging two other randomly selected replicates. If two of the four samples were missing for a given time point we imputed by taking the median expression of the remaining replicates. Accession O_302V was removed from the dataset due to missingness. We provide the code we used to normalize and filter in our RMarkdown (https://github.com/greenhamlab/B.rapa-diel-cold-regulation).

Before analyzing the transcriptomes we first assess each one by examining the variation among samples in a Principal Component Analysis and through hierarchical clustering. While the majority of the samples clustered relatively well (*i.e.* within a timepoint and/or treatment) we noticed one of FT005’s cold treated samples (Replicate 2; time point 6) appeared to be an outlier and was removed. We then imputed this sample as described above. To validate the RNA-sequencing we plot expression for a set of clock genes with known patterns and phases for each accession. Code for running and visualizing the PCAs, dendrograms, and expression patterns can also be found in our RMarkdown on GitHub.

### SNP-calling

Paired-end RNA-seq reads were trimmed using Trimmomatic (v0.33) and aligned to the *Brassica rapa* R500 reference genome (Brapassp_trilocularisR500_795_v2.0.fa) using STAR (v2.5.3a) in two-pass mode with read group information embedded in the BAM headers. PCR duplicates were marked using GATK MarkDuplicates (v4.1.2), and spliced alignments were processed with GATK SplitNCigarReads to remove intronic regions. Reads were filtered to retain only high-confidence alignments with mapping quality ≥20 using samtools (v1.21). BAM files from biological replicates of each accession were merged into a single BAM file using samtools.

Variants were jointly called across all accessions using GATK HaplotypeCaller (v4.1.2) in cohort mode with soft-clipped bases ignored. Jointly called variants were filtered using GATK hard-filtering following GATK best practices for RNA-seq variant calling. SNPs were filtered based on standard variant quality annotations, including quality by depth (QD), Fisher strand bias (FS), mapping quality (MQ), strand odds ratio (SOR), mapping quality rank sum (MQRankSum), and read position rank sum (ReadPosRankSum). Variants failing any of the following criteria were excluded: QD < 2.0, FS > 60.0, MQ < 40.0, SOR > 3.0, MQRankSum < −12.5, or ReadPosRankSum < −8.0. Only biallelic SNPs passing all quality filters were retained for downstream analyses.

Filtered SNPs were converted to PLINK format and pruned for linkage disequilibrium using a sliding window approach (50 kb window size, step size of 5 kb, and r² threshold of 0.2). Principal component analysis (PCA) was performed on the LD-pruned SNP set using PLINK (v1.90b6), and the first two principal components were visualized in R (v4.4.0) using ggplot2 to assess genetic relationships by morphotype among the 13 samples.

### Cold responsive transcriptomes

To identify genes with significantly different diel (24h) expression patterns after cold acclimation we used the time-series based R package DiPALM (Greenham et al. 2020). For the Weighted Gene Co-Expression Networks (WGCNA) (Langfelder and Horvath 2008) step, we estimated soft thresholding power and mean connectivity. We selected soft-thresholding powers for each accession by identifying the highest exponents (powers) that had reached stability with the lowest mean connectivity (Supplemental Data Set 2). Samples were collected from individual plants at each time point and assigned a replicate number. These replicates were then strung together to generate an expression profile for each gene. To account for inherent variation between these arbitrarily assigned plants we permuted DiPALM runs for each accession over 100 iterations and kept genes that had significantly different patterns (kME) or median expression (kMed) under cold in 95 of the 100 iterations. We use kME and kMed genes as our list of significantly cold responsive (CR) genes for the remainder of the analyses. These genes are provided as a zipped folder of csv’s for each genotype on our GitHub (https://github.com/greenhamlab/B.rapa-diel-cold-regulation/BrapaCRGeneLists.zip).

### Calculating Phase Change Groups (PCGs)

We took all genes with differential expression patterns (kMEs) and calculated a ‘Phase Change’ as the difference in max expression (phase) under control from the max expression (phase) under cold. This gave us the direction and magnitude of every phase change in four, eight, or 12 hour increments. Some differentially patterned genes had no change in phase but still exhibited differences in the overall pattern so they were included in the analysis. We collectively refer to these groups as ‘Phase Groups’.

Genes in each Phase Group are shifting in the same magnitude and direction, but they are not necessarily shifting at the same time of day relative to the control phase. To account for this, we further delineated the Phase Groups based on the time of day in control they were shifting from. This gave us 36 Phase Change Groups (PCGs) that encompass all 36 comparisons between the different magnitudes and directions (0; +/-4; +/-8; +/-12). The 12 hour changes are difficult to separate in terms of advancing or delaying, and as such should be interpreted accordingly. For our dataset we assigned advancing 12 hour PCGs first and then flipped the direction for the opposing group (*i.e.* ZT1 to ZT13 was assigned a +12 but ZT13 to ZT1 would be hard coded as a -12). To assess the biological functions enriched in our PCGs we ran Gene Ontology (GO) analyses using an in house script on each PCG and accession separately (Supplementary Data Set 3). We only kept terms that were significantly overenriched (Adjusted-p <= 0.05). See our RMarkdown for code to create the PCGs.

### Circadian Period by Leaf Movement

To determine the circadian period under control and cold acclimation conditions we used leaf-movement assays. We germinated seeds of each line on wet Whatman paper in petri dishes for two days and individual seedlings were transplanted into PVC pipe connectors filled with Jolly Gardener C/GP Germinating Mix soil. All plants were entrained for three days in 14h light/10h dark cycles with 22°C/16 °C temperature cycles. Control plants were released into free-running conditions at 22°C. To align the timing of leaf-movement with the RNA-seq dataset, cold-acclimated plants received an additional two days of entrainment under 14h light/10h dark, 10°C/6°C cycles followed by release into free-running conditions at 10°C. After 24h in free running conditions constant light and temperature, time-lapse imaging began. Images were captured every 20 minutes for one week using Canon Powershot 2000 cameras modified with the Canon Hack Development Kit (CHDK; https://chdk.fandom.com/wiki/CHDK_1.6_User_Manual). Image stacks were analyzed using Tracking Rhythms in Plants (TRiP) (Greenham et al. 2015), a computational pipeline that generates motion vectors for each plant and fits them to a circadian model to estimate circadian period length, which we have modified and uploaded to GitHub (https://github.com/GreenhamLab/TRiP). To visualize period length across lines by condition, data was plotted in R using the package ggplot2 (Villanueva and Chen 2019). A paired student’s t-test was conducted in R between cold acclimation and control conditions for each line using the RStatix package. Code for visualization and analyses are available in our RMarkdown on GitHub (https://github.com/greenhamlab/B.rapa-diel-cold-regulation).

### Gene Regulatory Networks

We built temporally informed Gene Regulatory Networks (GRNs) for control and cold data separately using the GENIE3 workflow (Huynh-Thu et al. 2010) for each accession. Known TFs from the Arabidopsis Plant Transcription Factor Data Base v.5.0 (https://planttfdb.gao-lab.org/index.php?sp=Ath) and known clock genes with known regulatory functions were used to make predictions. To determine a cutoff for significant edge weights we ran 100 permutations of each GRN for each accession and selected weights with a FDR cutoff of 0.05. To assess similarity among the networks we created a syntenic table anchored by the R500 genome (*Brassica rapa ssp. Trilocularis* v2.1) which allowed us to compare connections (also referred to as ‘edges’) across all accessions with different genomes. To assess conservation of predicted regulatory structure across accessions, we calculated pairwise Jaccard similarity of TF–target edge sets for all accession pairs within each treatment. Similarity was first calculated at the *B. rapa* gene level using accession-specific TF and target identifiers. To test whether retained-paralog divergence obscured shared regulatory architecture, *B. rapa* genes were then collapsed to orthology-aware identifiers based on their Arabidopsis orthologs (Supplemntary Data Set 7), and Jaccard similarity was recalculated using the ortholog-anchored edge sets. Comparisons were visualized using the R package pheatmap (R Core Team 2025). This process was repeated with the Arabidopsis ortholog to assess network similarity at the ortholog level. Data and scripts for this section can be found on our GitHub page (https://github.com/greenhamlab/B.rapa-diel-cold-regulation/GRNs).

### Shared cold-network construction

To identify regulatory connections retained across accessions with similar freezing responses, we constructed shared cold-network edge sets from the ortholog-anchored GRNs. Accessions were grouped based on relative freeze tolerance. The tolerant group included A03, VT123, PC185, and HN53; the sensitive group included R500, ZCT, and FT005; and the highly tolerant pair included PC185 and HN53. For each group, shared edges were defined using explicit support thresholds. Strict tolerant edges were present in all four tolerant accessions, relaxed tolerant edges were present in at least three of four tolerant accessions, strict sensitive edges were present in all three sensitive accessions, and relaxed sensitive edges were present in at least two of three sensitive accessions. PC185/HN53 edges were defined as those present in both accessions. These shared edge sets were used to identify group-supported TF–target relationships under cold acclimation.

### Integration of shared cold GRNs with phase-change GO themes

To connect shared regulatory structure with diel-retimed biological processes, we integrated the shared cold GRNs with phase-change group (PCG) GO enrichment results (Supplementary Data Set 3). Significant GO terms from accession-specific PCG enrichments were summarized into broad curated GO themes, including phosphorylation/kinase signaling, intracellular protein transport, vesicle trafficking/membrane coat, RNA binding/translation/ribosome, sulfur assimilation/redox homeostasis, and photosynthesis/light harvesting. For each candidate TF in the shared cold networks, we tested whether its predicted shared targets were enriched for genes assigned to each GO theme using Fisher’s exact tests. Candidate TF–theme relationships were retained for the Cytoscape visualization when the TF had at least three shared-network targets assigned to the GO theme, the TF-theme overlap passed Fisher FDR < 0.05, and the relationship was defined to the tolerant or sensitive edge class. Resulting TF–theme associations were classified by the shared-network support class from which they originated, including tolerant-associated, sensitive-associated, PC185/HN53-supported, or shared across tolerance groups. For Cytoscape visualization, cases in which the same TF was linked to the same GO theme through multiple support classes or sharing modes were collapsed to a single TF–theme edge. The collapsed edge retained the supporting evidence classes, sharing modes, best Fisher FDR, target-overlap counts, theme-coverage metrics, and overlapping target IDs. Target-level subnetworks were generated for selected GO themes by retaining TF–target edges connecting candidate TFs to GO-theme targets, allowing visualization of the inferred regulatory relationships underlying selected TF–theme associations.

### CNS Analysis

Conserved Noncoding Sequences were previously identified (Haudry et al. 2013; Supplementary Data Set 4) and filtered (Greenham et al. 2020) to generate a ∼65k set of CNSs found across crucifer genomes. Briefly, we followed previously published approaches (Greenham et al. 2020), using these sequences as BLAST queries (Camacho et al. 2009) against the newly-generated pan-*B. rapa* genomes (E-value cutoff of 0.01). We discarded all hits below 28.2 bitscore or 60% query coverage, and to account for the genome triplication in *B. rapa* since its ancestral divergence from Arabidopsis, we additionally discarded all high quality hits that aligned to more than three unique genomic loci. These candidate CNS sequences were then associated with their closest neighboring gene through bedtools closest (Quinlan and Hall 2010) and intersected with paralog, ortholog, and gene expression information. More detailed methods, as well as all scripts necessary to implement them, are available on the project GitHub page (https://github.com/greenhamlab/B.rapa-diel-cold-regulation).

## Results

### Genomic and freeze tolerance variation across a diverse *B. rapa* panel

To interpret accession-level variation in cold-acclimated transcriptomes, we first established the range of freeze tolerance across a panel of 14 *B. rapa* lines. These 14 accessions represented six morphotypes: vegetable turnip (VT123*, FT005), Chinese cabbage (HN53, CC168, A03*), choy sum (L58*), winter oil (OR213, WO83*), pak choi (PC185, ZCT, PCGlu*), and yellow sarson (R500*, ACC28, ACC50), allowing us to ask whether freezing response was primarily associated with morphotype or varied among accessions within morphotype. Plants were acclimated for 24h under chilling conditions and then exposed to a late-night freezing treatment at ZT17, a timing chosen to approximate the late-night to pre-dawn window when spring frost events commonly occur in temperate climates (Leske and Biddulph 2022) (Fig. 1a-b). We assessed freezing response using complementary physiological and performance traits, including visible leaf damage, electrolyte leakage, aboveground biomass, and light-adapted photosystem II efficiency (Fv’/Fm’) across the acclimation and recovery period (Fig. 1c-f; Fig. S1 and S2).

**Figure 1.**
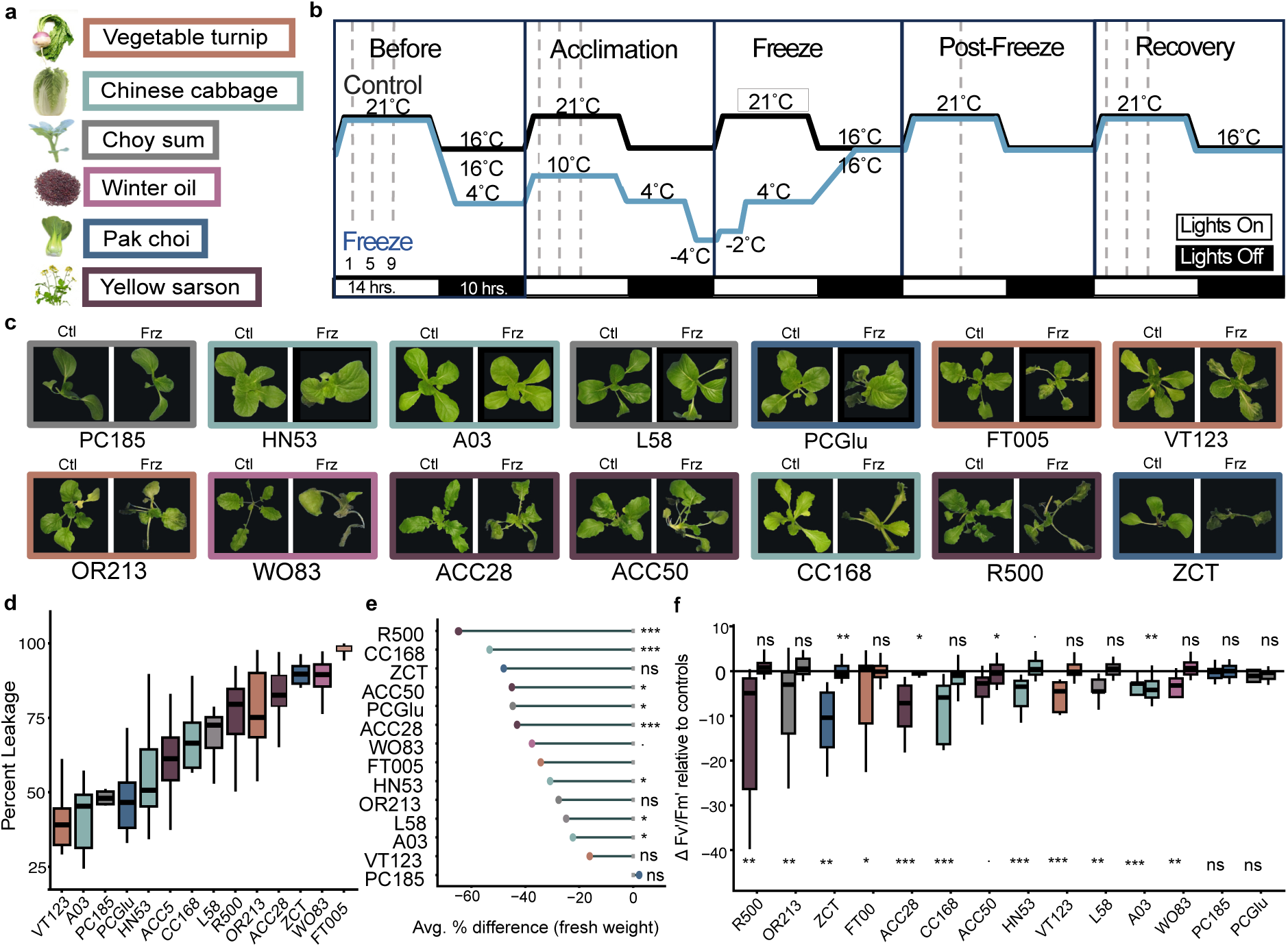
Relative freezing tolerance among six *Brassica rapa* morphotypes. (a) Six morphotypes were utilized in the freeze stress experiments which are color coded by morphotype. (b) Experimental design for all freeze stress assays. Freeze plants were cold acclimated for 24 hours followed by two hours of -4℃ and two hours at -2℃ before slowly warming back to control conditions. Dashed lines indicate Fv’/Fm’ collection at indicated ZTs (Zeitgeiber; hours after lights on). (c) Plant images are ordered by damage score with the lowest score on the top left and the highest damage score on the bottom right. Images were taken of freeze (Frz) and control (Ctl) the day following the freezing stress (‘Post-freeze’). Boxes are colored by their respective morphotype as described in panel A. (d) Electrolyte leakage assay quantifying the percent of ruptured cells following freezing stress. A lower percentage of leakage indicates a more tolerant plant. Boxes are colored by their respective morphotype. (e) Dots indicate the average reduction in above ground biomass following the freeze and are color coded by the respective morphotype. Two-sided Student t-Tests were run to determine significant differences in fresh weight. P-values for freeze-versus-control contrasts are indicated as follows: p ≤ 0.1 (.), p ≤ 0.05 (*), p ≤ 0.01 (**), and p≤ 0.001 (***). (f) Fv’/Fm’ measurements at ZT9 on the ‘Post Freeze’ day (left boxes) relative to recovery (right boxes). Percent recovery in Fv’/Fm’ was calculated as the difference between freeze stressed plants on day 4 (‘Post-Freeze’) and the difference on day 5 (‘Recovery’) from the mean of control responses taken on the respective day. Boxes are colored by morphotype, where lighter shaded boxes represent recovery responses (right). The line at 0 indicates no difference between freeze or recovery and the mean Fv’/Fm’ in control. Treatment effects were tested within each accession and sampling day using linear mixed-effects models with plant replicate as a random intercept to account for repeated measurements. P-values for freeze-versus-control contrasts were Holm-adjusted across all accession-by-day comparisons and are indicated as follows: adjusted p ≤ 0.1 (.), p ≤ 0.05 (*), p ≤ 0.01 (**), and p≤ 0.001 (***). Post-freeze treatment effects are shown along the bottom of the plot for each accession, and recovery treatment effects are shown along the top.

Freeze response varied widely across the panel and did not consistently follow morphotype. For example, all three yellow sarsons (R500, ACC50, and ACC28) had a moderate reduction in Fv’/Fm’ at ZT9 following the freeze, but ACC28 maintained less visible damage (Fig. 1c, S1) and smaller biomass reductions than R500 and ACC50 (Fig. 1e). Similarly, the two vegetable turnips, VT123 and FT005, had similar mean damage scores and drops in Fv’/Fm’ during acclimation (Fig. S1 and S2) yet FT005 had slower recovery of Fv’/Fm’ and high electrolyte leakage while VT123 had the lowest leakage of all accessions (Fig. 1d, 1f). The three Chinese cabbage accessions (CC168, A03, and HN53) were more similar in electrolyte leakage and Fv’/Fm’; although CC168 showed larger reduction in aboveground biomass than HN53 or A03 (Fig 1d-f). Across all metrics, PC185 (pak choi) and HN53 (chinese cabbage) consistently showed the strongest freeze tolerance, whereas R500 (yellow sarson) and ZCT (pak choi) were the most sensitive, followed by FT005 (vegetable turnip). Thus, the panel captures substantial accession-level variation in freezing response, including variation within morphotypes, providing a phenotypic framework for comparing diel cold-acclimated transcriptomes.

To interpret physiological differences (Fig. 1) in the context of accession-specific gene content and regulatory sequence variation, we generated a new *B. rapa* pangenome reference for six representative morphotypes (* lines above), each coassembled with deep HiFi (coverage = 41-126X) and Omni-C (72-182X), polished with Illumina 2×150bp reads (50-65X, Supplementary Data Set 1). The 2,965Mb of nearly-gapless genomic DNA sequence across the six assemblies was highly collinear, containing only a handful of large structural variants (‘SVs’) including only 12.4Mb across 7 unique inversions larger than 1Mb (Fig. 2a). It is noteworthy that SVs were far more common when called against the original V1 R500 genome (Lou et al. 2020). Relative to our new R500 “V2” assembly, V1 contained many large gaps and assembly errors that have now been resolved (Fig. S3).

**Figure 2.**
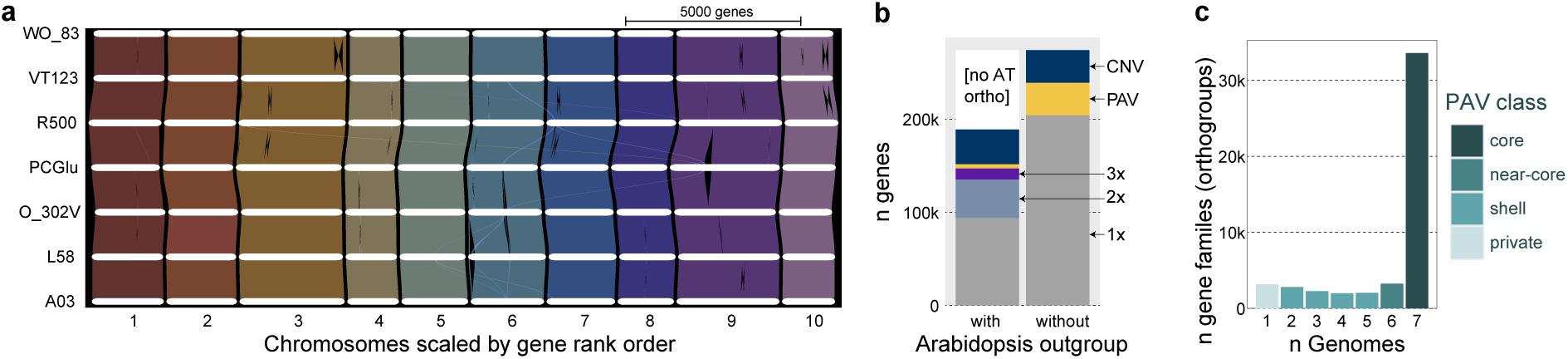
Structure of pangenomic variation. (a) Synteny across the seven genomes visualized as collinear blocks of genes via GENESPACE. (b) Gene copy number variation across the pan-gene set, calculated as membership in Orthofinder’s phylogenetically hierarchical orthogroups. Counts are split by orthology inferred with and without the Arabidopsis araport 11 genome annotation in the species tree. (c) Distribution of gene presence absence variation across the seven annotations.

We complemented the genome assemblies with thorough and high-confidence protein-coding gene annotations (Supplementary Data Set 1). The resulting pangene set includes 274,016 gene models and 44,796 orthologous sets of genes (‘orthogroups’). By considering *Arabidopsis thaliana* as an outgroup, we were also able to frame orthogroups in the context of the ancient Brassicaceae whole-genome triplication (“WGT”) (Cheng et al. 2014). Among the 20,855 orthogroups that contained *A. thaliana* genes, 570 and 2975 were uniformly 2- and 3-copies all *B. rapa* genomes: 17.0% of all gene families retained at least one paralog from the WGT in all *B. rapa* genomes (Fig. 2b). However, 14,413 orthogroups (69.1%) were uniformly single copy in all *B. rapa* genomes and we observed substantial presence-absence variation across the pangene set (Fig. 2c). Combined, these results demonstrate that while *B. rapa* genomes are well on their way towards “diploidization”, many families retain duplicates, which could have evolved away from redundancy via processes of neo- and sub-functionalization and provide the raw genetic material for evolution of stress responses and other adaptive traits (Panchy et al. 2016; Soltis and Soltis 2016; Almeida-Silva and Van de Peer 2023; Edger et al. 2026).

### Diel transcriptome re-wiring during cold acclimation

To determine how cold acclimation reshapes gene expression across the day, we generated a diel RNA-seq time course across the *B. rapa* accession panel. Because stress responses, photosynthesis, metabolism, and growth are strongly structured by time of day (Wilkins et al. 2009; Graf et al. 2010; Dodd et al. 2015; Lu et al. 2021; Bonnot et al. 2023; de Barros Dantas et al. 2023), we sampled leaf tissue every four hours over a 24h period after three days of cold acclimation. This design allowed us to distinguish genes that changed overall expression under cold from genes whose timing across the diel cycle was altered (Fig. S3). Before comparing transcriptional responses, we asked whether freezing tolerance was primarily associated with genetic background. Genome-wide single-nucleotide polymorphism (SNP) analysis showed that accessions clustered largely by morphotype rather than by relative freeze tolerance, indicating that the tolerance and sensitive phenotypes observed in the freeze assays could not be explained simply by broad population structure (Fig. S4). This supported a transcriptome-wide comparison focused on accession-specific cold response and time-of-day regulation.

Cold acclimation caused extensive transcriptome reprogramming across the panel. Using DiPALM (Greenham et al. 2020), we identified genes with altered median expression and genes with altered expression patterns across the diel time course in each accession. Each accession contained thousands of cold-responsive genes, indicating broad transcriptional remodeling under acclimation conditions (Supplemental Data Set 2). For the differentially patterned genes, we calculated the difference between the time of peak expression (phase) under cold and control conditions, assigning each gene to a phase-change category (Fig. 3A). We left out PCGlu for this analysis as nearly all of the cold-responsive genes exhibited changes in median expression instead of a pattern change. Across accessions, cold-responsive genes spanned a wide range of phase changes, including ±4h, ±8h, ±12h and altered patterns with no change in phase (0). Large changes of ±12 were relatively uncommon, whereas smaller phase changes were more frequent across the panel.

**Figure 3.**
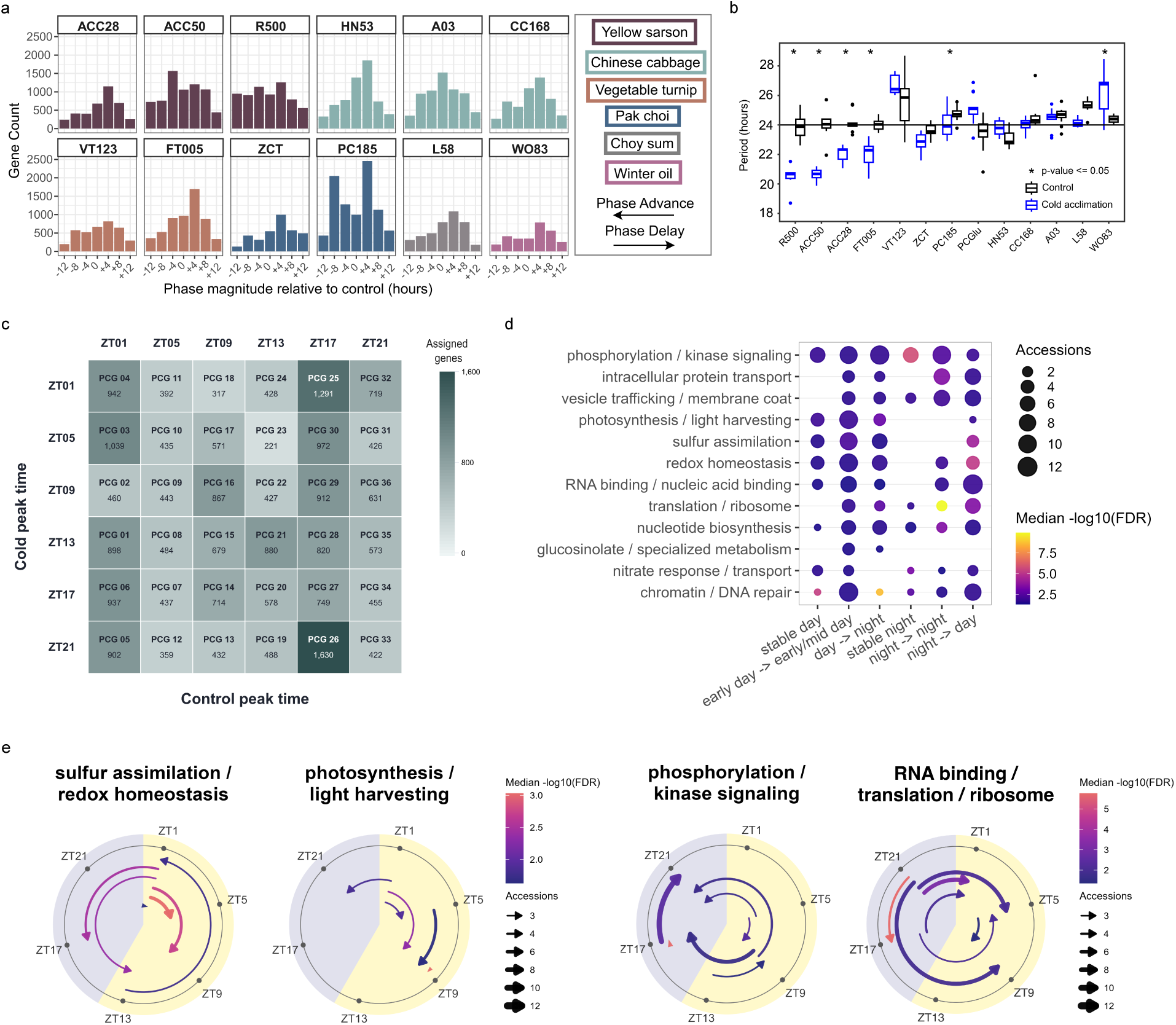
*B. rapa* accessions display distinct cold-induced phase changes and variation in endogenous circadian period length. (a) Phase changes were calculated as the difference between the max expression in control (phase) from the max expression in cold. Phase changes are along the x-axis and are measured in hours. The “0” group contains genes that had a significantly different pattern but did not have a change in phase between treatments. Morphotypes are colored based on the key on the right and accessions are labeled at the top of each facet. (b) Period length in hours as determined by leaf-movement in constant light and temperature for Cold (12°C) and Control (22°C) for each accession. A Student’s t-Test was conducted between control (black) and cold (blue) acclimation groups. The line represents period values at 24h as expected under diel conditions. Statistically significant responses are denoted with an asterisk (p-value <0.05). accession identifiers are listed at the bottom. (c) Phase Change Group (PCGs) designations with hours after lights on (ZTs) during control (top) and cold treatments (left) with the PCG identifier in each cell. Numbers within the cells indicate the number of genes assigned to a given PCG. Darker colors indicate more genes in a given PCG. (d) Enrichment of curated GO themes (y axis) and the general time of day these themes shift to under cold acclimation. The size of each bubble represents the number of accessions with enrichment in a given theme and the colors describe significance where warmer colors indicate a smaller FDR (False Discovery Rate). Temporal shifts are described along the x-axis. Stable day/stable night groups are still cold responsive but did not exhibit a change in phase. (e) All phase changes for broader GO themes identified in (d). Samples were collected every four hours for 24h as indicated around the outside of each circle. Daytime phases are highlighted in yellow and nighttime samples in blue. Arrows are colored by the median FDR for PCGs of accessions that had enrichment in the given pathway, and flow from the control phase (flat end) to the shifted phase under cold acclimation (arrowhead).

The distribution of phase changes varied strongly by accession and did not follow a simple morphotype or tolerance-class pattern (Fig. 3A), although in general the largest proportions were found in the ±4h groups. For example, PC185, one of the most freeze-tolerant accessions, had a relatively large portion of cold-responsive pattern change genes in the +8h and -4h groups, whereas HN53, another tolerant accession, had most cold-responsive pattern change genes in 0 and +4h groups. Thus, even accession with similar freeze tolerance can differ in how cold alters the timing of gene expression. These results indicate that cold acclimation produces accession-specific diel transcriptome rewiring rather than a uniform temporal response across *B. rapa*.

Given the large portion of the *B. rapa* transcriptome under circadian control (Greenham et al. 2020), one explanation for the variation in phase changes across accessions is differences in circadian period in combination with altered temperature compensation, the ability to maintain a ∼24h circadian period across physiologically-relevant temperatures (Maeda and Nakamichi 2025). We next asked whether accession-specific differences in circadian timing could contribute to the observed phase changes. Circadian period was determined by leaf movement (Greenham et al. 2015) under both control and cold entrainment conditions. Leaf movement assays revealed variation in endogenous period under both control and cold conditions (Fig. 3B). Under control conditions, most accessions had circadian periods close to 24h, although several accessions showed longer and shorter periods, including VT123 (25.5h), L58 (25.3h), HN53 (23h), and PCGlu (23.5h). Under cold conditions, some accessions were relatively well compensated, including PC185, L58, and all three chinese cabbages HN53, CC168, and A03. In contrast, R500, ACC50, ACC28, FT005, and ZCT showed shortened periods under cold, whereas WO83, PCGlu and VT123 showed longer periods. These differences in period and temperature compensation could contribute to the accession-specific phase changes observed in the RNA-seq time course.

Because the transcriptome experiment was conducted under diel light/dark and temperature cycles, we cannot fully separate direct cold responses from changes in circadian phase, temperature compensation, or other diel environmental responses. However, this interaction is biologically relevant since under natural cold acclimation, plants experience low temperature in the context of daily changes in light and temperature. Together, the RNA-seq and leaf movement data indicate that cold acclimation in *B. rapa* is strongly time structured and that accession-specific differences in circadian timing may contribute to how cold-responsive genes are redistributed across the diel cycle.

### Phase-change groups reveal recurrent diel-retiming of core biological processes

To identify biological processes associated with cold-induced changes in diel expression timing, we grouped pattern change genes by their control-to-cold phase transition, generating 36 Phase Change Groups (PCGs; Fig. 3c). We then performed GO enrichment separately for each accession and PCG. Because GO terms are often redundant and difficult to compare across many accession-by-PCG combinations, we summarized significant terms into broad, manually curated process themes (Fig. 3c; Supplemental Data Set 3). These themes were used as a hypothesis-generating framework to identify recurrent biological patterns across PCGs.

Several process themes recurred across accessions, but the timing of those enrichments often differed indicating that cold causes some pathways to remain in the same diel window while others were reassigned from day to night, night to day, or shifted within day/night. Phosphorylation/kinase signaling was the most broadly represented theme, with 33 GO terms enriched across all 13 accessions, 27 PCG transitions, and 891 contributing *B. rapa* genes (Supplemental Data Set 3). Prominent transitions included an early-night to late-night shift (ZT17 to ZT21) and a midday to early-night shift (ZT9 to ZT17), indicating that phosphorylation-related genes were repeatedly associated with changes in diel timing under cold (Fig. 3d,e). Intracellular protein transport and vesicle trafficking/membrane coat showed related nighttime-shifted patterns (Fig. 3d), suggesting that signaling, protein movement, and vesicle-associated processes are recurrently retimed during cold acclimation.

RNA binding/translation/ribosome-related terms formed the strongest theme statistically, with 29 GO terms enriched across 12 accessions, 18 PCG transitions, and 1,108 contributing *B. rapa* genes. This theme was particularly prominent across night-to-day and stable-night transition classes (Fig. 3d,e), suggesting that cold acclimation may alter the timing of RNA-associated processes and translational capacity across the diel cycle. Photosynthesis/light harvesting formed a smaller but biologically coherent theme, with 20 GO terms enriched across nine accessions, 16 PCG transitions, and 160 contributing *B. rapa* genes. Many of these enrichments involved shifts within the day or toward later daytime expression under cold (Fig. 3d,e), consistent with altered timing of light-harvesting and photosystem-related processes during acclimation.

Sulfur assimilation and redox homeostasis were enriched in the majority of accessions and across several timing transitions. Sulfur assimilation/redox homeostasis showed a distinct pattern that was not as large as the phosphorylation or RNA/translation themes, but was broadly recurrent and temporally striking. This theme included 24 GO terms enriched across all 13 accessions, 20 PCG transitions, and 108 contributing *B. rapa* genes. In contrast to phosphorylation and translation, sulfur/redox-associated enrichments often originated from genes peaking near dawn under control conditions, but these genes shifted to multiple cold-phase destinations across accessions (Fig. 3e). This suggests that sulfur assimilation and redox-associated processes may be similarly timed under non-stress conditions but become reallocated to different diel windows during cold acclimation in an accession-specific manner. Given the roles of sulfur metabolism and redox buffering in stress physiology (Foyer and Noctor 2005; Hossain et al. 2017; Sun et al. 2026; Wang et al. 2026), this pattern points to sulfur/redox timing as a potential axis of accession-specific acclimation strategy.

Although these themes recurred across the panel, the specific PCG transitions and accessions contributing to each theme differed substantially. Thus, cold acclimation does not appear to impose a single shared temporal program across *B. rapa* accessions. Instead, similar biological processes are retimed in different ways across accessions. Additional curated themes, including glucosinolate/specialized metabolism, nitrate response/transport, chromatin/DNA repair, nucleotide biosynthesis, and hormone signaling, were enriched in subsets of accessions rather than forming the primary recurrent modules emphasized here. Together, these PCG/GO patterns indicate that cold acclimation in *B. rapa* involves both recurrent retiming of core cellular processes and accession-specific temporal reorganization of signaling, translation, photosynthesis, sulfur/redox metabolism, nutrient regulation, and specialized metabolism.

### Ortholog-anchored GRNs reveal conserved regulatory architecture obscured by paralog diversification

To identify transcriptional regulators associated with accession-specific temporal responses to cold, we inferred control and cold Gene Regulatory Networks (GRNs) for each of the *B. rapa* accessions using GENIE3 (Huynh-Thu et al. 2010). We first compared networks at the *B. rapa* gene level by calculating pairwise Jaccard similarity of predicted TF-target edges across accessions. Overall, similarity among control networks was low, indicating substantial divergence in predicted regulatory connections among *B. rapa* accessions (Fig. 4a). A small number of accession pairs showed higher similarity, including the relatively intolerant yellow sarsons R500 and ACC50 and the Chinese cabbage CC168 with vegetable turnip VT123 and pak choy ZCT (Fig. 4a). In contrast, PC185 and A03 shared relatively few predicted edges with other accessions. Under cold acclimation, pairwise similarity was further reduced across all comparisons, consistent with accession-specific regulatory rewiring in response to cold. In contrast to the control, the highest similarity was found between the Chinese cabbage A03 and HN53 followed by the two yellow sarsons R500 and ACC50 (Fig. 4a).

**Figure 4.**
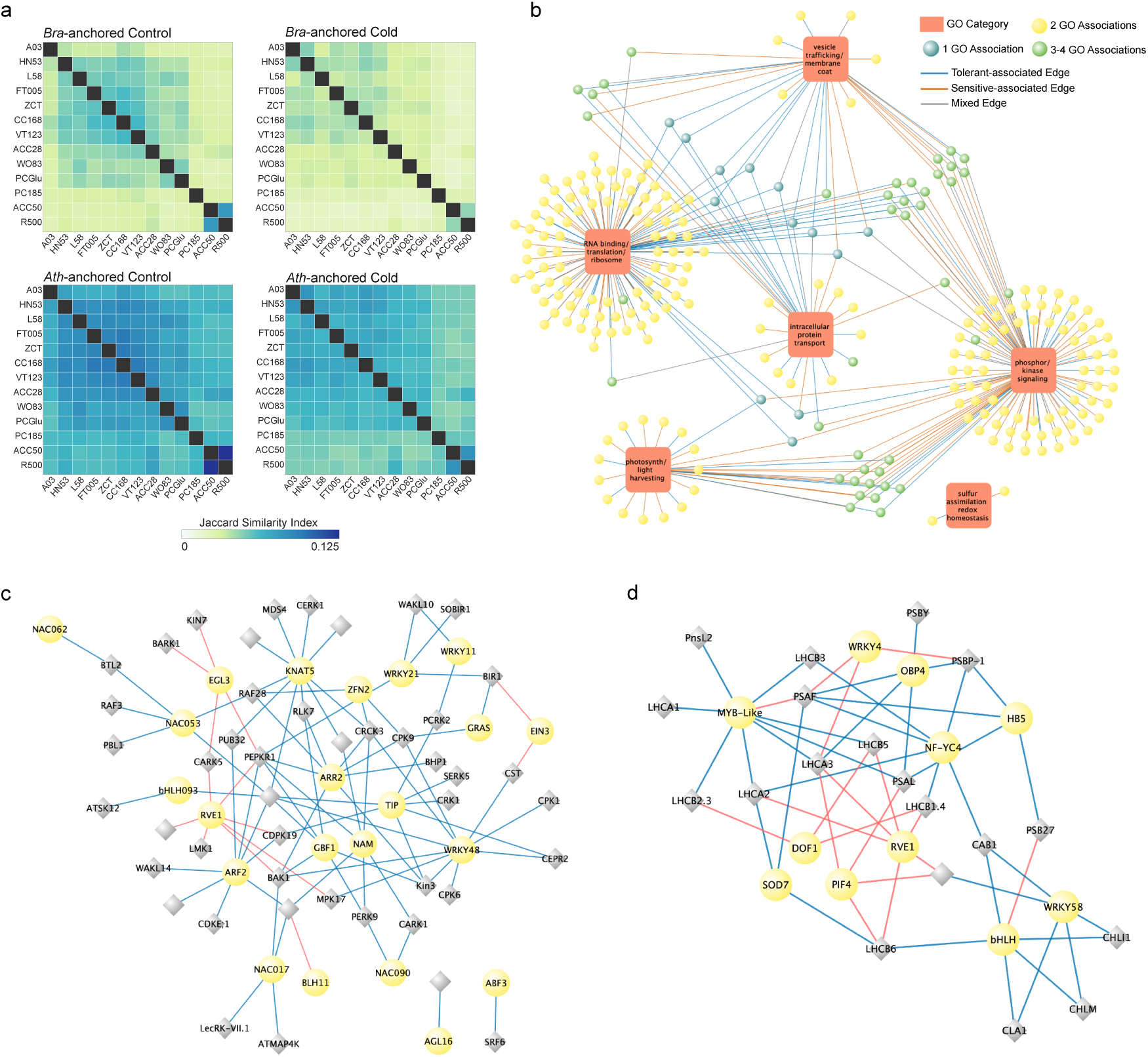
Cytoscape network showing candidate transcription factors associated with six recurrent GO themes identified from cold-induced diel phase-change groups. (a) Jaccard similarity indices where darker colors indicate more similar network edges among accessions. (b) Orange rounded rectangles represent GO-theme nodes, and circular nodes represent candidate TFs. Edges connect TFs to GO themes when the TF’s predicted shared cold-network targets significantly overlapped genes assigned to that GO theme. TF node color indicates the number of GO themes associated with each TF, with yellow indicating one theme, light green indicating two themes, and teal indicating three to four themes. Edge color indicates the shared-network evidence class for the TF–theme relationship, distinguishing tolerant-associated (blue), sensitive-associated (orange), and cross-group shared regulatory associations (grey). For clarity, duplicate TF–GO-theme edges were collapsed to one edge per TF–theme pair while retaining combined evidence attributes. (c) Cytoscape subnetwork showing candidate TF–target relationships within the phosphorylation/kinase signaling GO theme. This network was filtered for strict tolerant-associated edges connected to targets assigned to the ZT17→ZT21 phase-change group. Yellow circular nodes represent candidate TFs, and gray diamond nodes represent phosphorylation-associated target genes. Blue edges indicate TF–target relationships detected in the PC185/HN53 shared network or shared between the tolerant group and PC185/HN53, whereas red edges indicate relationships detected only in the tolerant shared network. (d) Cytoscape subnetwork showing candidate TF–target relationships within the photosynthesis/light-harvesting GO theme. This network was filtered for strict tolerant-associated edges. Color scheme is as described in (C).

One explanation for the low conservation of *B. rapa* gene-level GRNs is that shared regulatory pathways may be obscured by divergence among retained paralogs. Previous work in *B. rapa* showed that retained *B. rapa* paralogs frequently diverge in temporal expression patterns under circadian and diel conditions (Greenham et al. 2020), suggesting that accessions may deploy different paralogs within otherwise related regulatory pathways. To test whether paralog-level divergence contributed to the low GRN similarity, we collapsed *B. rapa* genes to their Arabidopsis orthologs and recalculated the pairwise network similarity. This ortholog-anchored comparison substantially increased network similarity across accession pairs in both the control and cold conditions (Fig. 4a). Thus, although predicted regulatory edges are highly accession-specific at the retained-paralog level, a stronger layer of shared regulatory architecture emerges when networks are compared at the orthologous pathway level. This result indicates that part of the apparent network divergence among *B. rapa* accessions reflects paralog-level diversification rather than complete loss of conserved regulatory programs. Despite the increase in ortholog-level conservation, cold-acclimated networks still showed reduced similarity relative to control networks, indicating that cold induces additional accession-specific rewiring even when paralog identity is collapsed. Together, these results support a model in which conserved regulatory pathways are implemented through divergent retained paralogs across *B. rapa* accessions, and cold acclimation further reshapes these networks in an accession-specific manner.

### Shared cold GRNs connect candidate TFs to diel-retimed biological processes

Having established that ortholog anchoring reveals conserved regulatory architecture across accessions, we next asked whether these retained cold-network connections were associated with the biological processes that showed the recurrent diel retiming under cold acclimation. To test this, we integrated the PCG/GO enrichment results with the cold GRNs by linking candidate TFs to GO themes when their predicted shared cold-network targets significantly overlapped genes assigned to those themes. This analysis identified 314 TF-theme associations involving 250 candidate TFs across the six recurrent GO themes (Fig. 4b). The strongest convergence occurred around RNA binding/translation/ribosome and phosphorylation/kinase signaling, which accounted for 114 and 111 TF–theme associations, respectively. In contrast, photosynthesis/light harvesting, vesicle trafficking/membrane coat, and intracellular protein transport formed smaller modules with 30, 29, and 28 TF–theme associations, respectively, while sulfur assimilation/redox homeostasis was represented by two TF–theme associations. Thus, the shared cold networks were not randomly distributed across PCG/GO themes, but were concentrated around a subset of recurrent diel-retimed processes.

We next asked whether candidate TFs were generally associated with single processes or connected to multiple diel-retimed themes. Most TFs were associated with one GO theme (200 out of 250), whereas 37 TFs were linked to two themes and 13 TFs were linked to three or four themes (Fig. 4b). These multi-theme TFs may represent candidate regulators with broader roles in coordinating the diel cold response, although the majority of inferred regulators appeared process-specific within this analysis. Each TF-theme association was classified according to whether the underlying shared cold network edges were supported by tolerant accessions (A03, VT123, PC185, and HN53), sensitive accessions (R500, ZCT, and FT005) or both groups. The presence of TF-theme associations across all three edge classes suggests that some cold-retimed biological processes are associated with regulatory structure shared across tolerance groups, whereas others include candidate TF-theme relationships preferentially retained in tolerant or sensitive shared networks.

To move from broad TF-theme associations to the underlying inferred regulatory edges, we focused on two representative tolerant-associated subnetworks. First, because phosphorylation/kinase signaling was one of the most recurrent diel-retimed themes and showed a prominent ZT17->ZT21 transition, we filtered this theme for strict tolerant-associated TF-target edges connected to ZT17->ZT21 targets. This produced a compact regulatory network of 99 TF-target edges connecting 25 candidate TFs to 48 phosphorylation-related targets (Fig. 4c). The network included several highly connected candidate regulators such as RESPONSE REGULATOR2 (ARR2), KNOTTED-LIKE FROM ARABIDOPSIS THALIANA 5 (KNAT5), AUXIN RESPONSE FACTOR 2 (ARF2), REVEILLE 1 (RVE1), NAC domain containing protein 17 (NAC017), and NAC052. The target set included receptor-like kinases, calcium-dependent protein kinases, MAP kinase-related genes, and other phosphorylation-associated components (Fig. 4c). Most edges in this network were supported by PC185/HN53 or shared across tolerant accessions (Fig. 4c). These results identify tolerant-associated inferred regulatory relationships linked to a nighttime-retimed phosphorylation/kinase signaling module.

We next examined photosynthesis/light harvesting because it formed a smaller and more discrete module in the TF-to-GO network and was enriched among PCGs involving shifts from morning or early-day expression toward later daytime expression under cold. The photosynthesis subnetwork contained 54 tolerant-associated TF-target edges connecting 11 candidate TFs to 19 photosynthesis associated targets (Fig. 4D). These targets included *PHOTOSYSTEM I LIGHT HARVESTING COMPLEX* genes *LHCA1, LHCA2,* and *LHCA3* as well as *LIGHT HARVESTING COMPLEX OF PHOTOSYSTEM II* genes *LHCB1.4*, *LHCB*2*.3*, *LHCB3*, *LHCB5*, and *LHCB6* (Fig. 4D). Several candidate regulators including NUCLEAR FACTOR Y (NF-YC4), WRKY4, PHYTOCHROME INTERACTING FACTOR 4 (PIF4), and RVE1 were connected to multiple photosynthesis-associated targets, with RVE1 showing tolerant-shared connectivity in this module. Together, the phosphorylation and photosynthesis subnetworks suggest that tolerant accessions retain specific inferred TF–target relationships associated with diel-retimed signaling and photosynthetic processes. These networks do not establish direct regulation, but they prioritize candidate regulators and target modules for testing how altered timing contributes to cold acclimation.

### Conserved Non-coding Sequences (CNSs) across the *B. rapa* pangenome highlight cases of paralog family diversification

The ortholog-anchored GRN analysis suggested that conserved regulatory programs can be obscured by retained-paralog divergence. We next asked whether conserved noncoding sequences (CNSs) could identify candidate cis-regulatory features associated with this paralog-specific diversification. Using previously described CNSs across the Brassicaceae (Haudry et al. 2013; Supplemental Data Set 4), we mapped CNSs across the six *B. rapa* pangenome accessions and examined how observed highly specific temporal retiming trends might have evolved (Supplemental Data Set 5). As expected for conserved regulatory elements, CNS-anchored gene relationships were highly similar across the accession-resolved assemblies (Fig 5a). At the individual gene level, we observed variable ortholog-CNS sharing depending on the filtering criteria. Across all identified ortholog pairs, approximately 33% shared a CNS; however, when the comparison was restricted to genes where both the gene and its ortholog had at least one associated CNS, the ortholog partner shared that same CNS approximately 80% of the time (Fig 5b). Thus, when CNSs are detected in both orthologous regions, their assignments are usually preserved across accessions.

**Figure 5.**
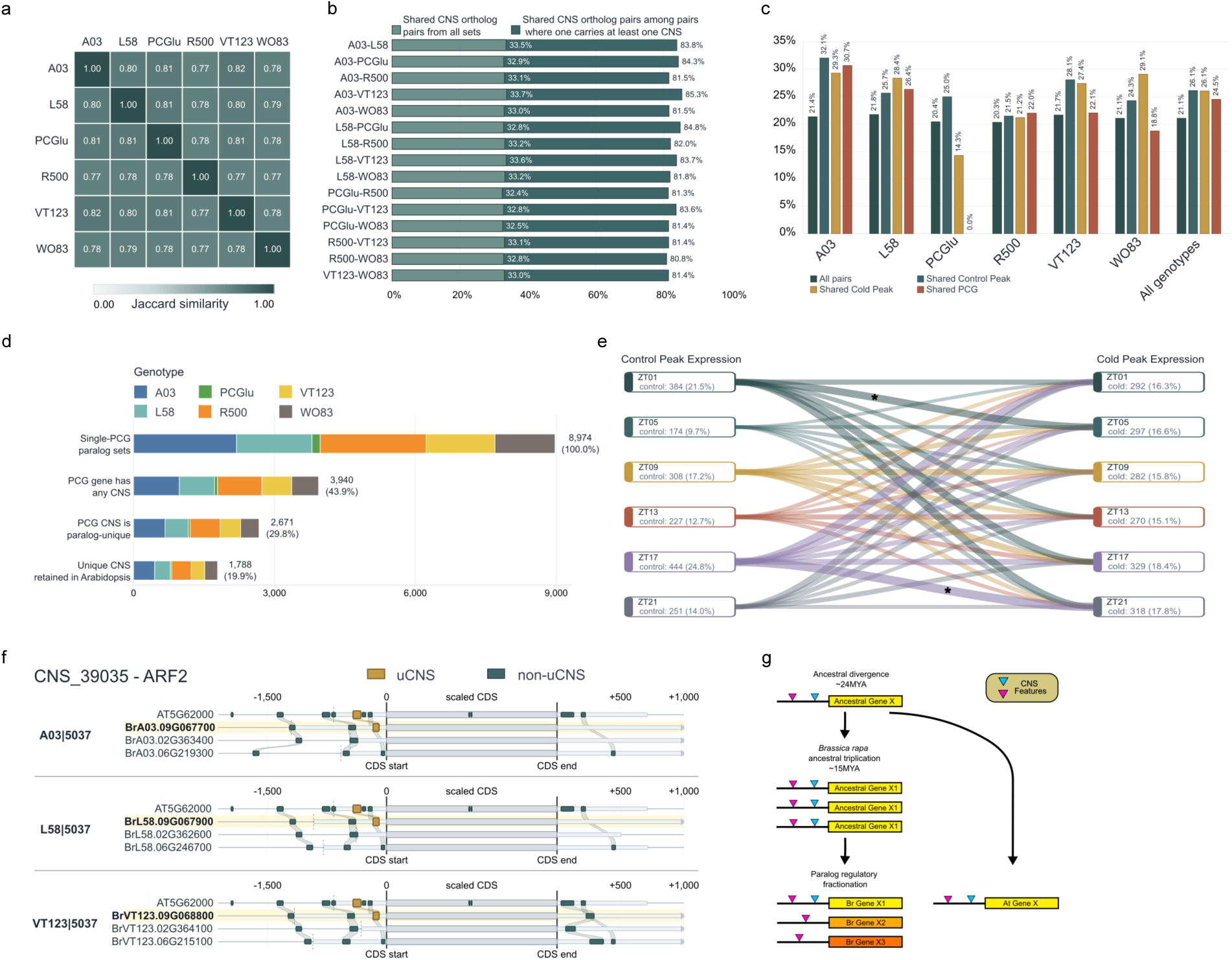
Divergence in a Conserved Non-coding Sequences in *ARF2* paralogs assigned to disparate Phase Change Groups. (a) Jaccard similarity scores using previously identified Conserved Non-coding Sequences (CNSs) mapped to each morphotype. (b) Percentage of all syntenic ortholog genes that share a CNS (lighter hue), and the same proportion when requiring CNS presence in at least one of the two orthologs. (c) Conserved CNSs among paralogs that are either assigned to the same Phase Change Group (PCG; red), that have the same phase under control (dark blue), the same phase under cold (yellow), or have the same CNS among all paralog pairs (black). (d) Number of paralog sets across all genotypes where a single member is uniquely assigned to a PCG (top bar), further filtered to require that the unique PCG-assigned gene also contains at least one CNS (second bar), and that that one CNS is unique among the paralog family (third bar). The bottom bar then identifies which of those paralog sets with a unique PCG-assigned genes associated with a unique CNS also share that unique CNS with their Arabidopsis ortholog. (e) Phase changes between control and cold for paralog sets that have retained a ‘unique’ CNS as identified in (d). Numbers of genes that meet this criteria across all accessions, and proportions of genes relative to total expressed genes, are listed below the time of day (ZT) identifier in each box. Ribbon sizes depict the number of genes in that group and are colored by their control phase. Asterisks indicate shifts with the largest number of genes with altered timing under cold. (f) Gene models highlighting divergence in a unique CNS (uCNS; yellow squares) among paralogous *B. rapa* copies in the *ARF2* Arabidopsis ortholog. Gene models are stacked by accession as indicated on the left. Non-unique CNSs (non-uCNS) are indicated by the dark blue squares. Ribbons show relative positions between CNS content among orthologs. Coding sequences are highlighted in grey and scaled to a uniform length. (G) Evolutionary model depicting divergence of regulatory sequences among paralogs following the *B. rapa* triplication event. Following triplication, a CNS (blue) is fractionated and lost in two of the three copies, leading to a single ortholog with the ancestral CNS.

Retained paralogs showed a markedly different pattern. Within each accession, paralog pairs shared a CNS only approximately 21% of the time (Fig 5C), indicating substantially greater divergence in CNS architecture among paralogs than among orthologs. Generally, CNS sharing was higher among paralogs assigned to the same control peak expression, cold peak expression, or phase change group (Fig 5c), suggesting that shared CNS architecture is associated with more similar diel expression behavior. PCGlu is one exception due to it having very few pattern change genes. However, the majority of paralogs did not share the same CNSs, consistent with extensive cis-regulatory diversification after genome triplication.

We chose to focus on a subset of paralogs that could link CNS divergence to cold-induced diel retiming. First, we identified ∼9000 paralog families across the six CNS-annotated *B. rapa* genomes where only a single member in each paralog family was assigned to a PCG, and noted that ∼4000 of these families contained at least one CNS association (Fig 5d). Of these genes, ∼2700 paralog families were structured such that the sole-PCG-assigned member was also associated with a unique CNS (uCNS) not identified elsewhere in the paralog set (Fig 5d). These paralog-CNS relationships could arise through several evolutionary routes, including gain of a novel CNS or loss of an ancestral element after the *B. rapa* genome triplication. To distinguish candidates consistent with ancestral CNS retention, we asked whether the same CNS was present near the Arabidopsis ortholog, which diverged from the *B. rapa* lineage before genome triplication (Cheng et al. 2014). This identified approximately 1,800 genes in which the sole-PCG-assigned gene within a paralog set uniquely shared a CNS (uCNS) with its Arabidopsis ortholog (Fig 5d, Supplementary Data Set 6). These genes represent strong candidates where an ancestral cis-regulatory element may have been retained near one paralog while being absent from other retained paralogs.

The ∼1800 Arabidopsis-matched, paralog-uCNS genes were distributed across control peak phases, cold peak phases, and PCGs (Fig 5e), indicating that this pattern is not restricted to a single expression class. The largest number of control peak assigned genes occurred at ZT17, while cold peak assigned genes were more evenly distributed. Among PCGs, while all retiming categories were represented, the ZT17 -> ZT21 shift was the largest at 118 genes, followed by the ZT01 -> ZT05 shift containing 91 genes. The enrichment of candidates in the ZT17->ZT21 group is notable because this same transition was prominent in the phosphorylation/kinase signaling GO theme and in the tolerant-associated phosphorylation GRN subnetwork. Thus, the CNS analysis converges with the PCG/GO and GRN analyses on a nighttime-retimed regulatory context.

The *ARF2*-associated CNS_39035 illustrates this pattern at a single locus (Fig 5f). This uCNS is associated with the Arabidopsis *ARF2* ortholog, and with one member of the *B. rapa* paralog sets in A03, L58, and VT123 (Fig 5f, yellow boxes). In all three accessions, this CNS is present near the coding sequence in one of the three copies, but missing in the other two; however, additional adjacent and nearby CNSs map cleanly between the *B. rapa* accessions and across to Arabidopsis. This arrangement is consistent with a model in which ancestral regulatory elements were retained across the Arabidopsis-*B. rapa* split, but underwent differential fractionation among *B. rapa* paralogs after genome triplication, ultimately generating the paralog-specific combinations of uCNSs and CNSs observed here (Fig 5g). Although these data do not demonstrate that the CNS_39035 directly controls *ARF2* expression timing, they show how pangenome-enabled CNS mapping can prioritize paralog-specific regulatory elements associated with cold-induced diel retiming.

## Discussion

Improving crop tolerance to temperature extremes requires understanding not only which genes respond to stress, but when those responses occur. Stress acclimation must be coordinated with photosynthesis, growth, nutrient assimilation, and metabolism (Huot et al. 2014), all of which are strongly structured by time of day (Dodd et al. 2005). Here, we used a panel of *B. rapa* accessions with diverse freeze tolerances, alongside a newly generated pangenome spanning six morphotypes, to examine how cold acclimation reshapes diel transcriptome regulation. We found that cold acclimation extensively reprograms gene expression across the diel cycle, but the timing and regulatory architecture of these responses vary substantially among accessions. This is consistent with previous work in *B. rapa* (Greenham et al. 2017, 2020), poplar (Robertson et al. 2022), and Arabidopsis (Wilkins et al. 2010) which show that the largest variation in stress responses is attributed to time of day. Together, our results support a model in which cold acclimation in *B. rapa* involves recurrent retiming of core biological processes, accession-specific deployment of retained paralogs, and differential retention of regulatory connections that may contribute to freeze tolerance.

A major finding of this study is that freeze tolerance varied significantly within morphotypes. Although accessions clustered by morphotype at the genome-wide SNP level, physiological responses to freezing did not follow morphotype alone. For example, PC185 and HN53 consistently showed relatively strong freeze tolerance across physiological metrics, whereas R500, ZCT, and FT005 were among the most sensitive. This within-morphotype variation suggests that cold tolerance is not simply explained by broad population structure or crop type. Instead, tolerance likely reflects accession-specific combinations of physiological and regulatory traits. This variation provided an important framework for interpreting the transcriptomic and network analysis, allowing us to ask whether tolerant accessions share regulatory features that are not apparent from morphotype or genetic background alone.

Given that a substantial proportion of plant transcriptomes is under circadian regulation (Covington et al. 2008; Greenham et al. 2020; Ricono et al. 2025), including stress-related pathways (Greenham and McClung 2015; Sanchez and Kay 2016), we hypothesized that shifts in the timing, rather than only changes in expression magnitude, would represent a major source of regulatory divergence among accessions. Our time-of-day-resolved transcriptomics revealed that cold acclimation extensively altered thousands of diel expression patterns across the panel. Rather than producing a uniform transcriptional response, cold shifted the timing of peak expression for many genes, with accessions differing in both the number of cold-responsive genes and the distribution of phase-change groups. These findings are consistent with other studies showing that stress-responsive expression cannot be fully understood from single time-point comparisons (Greenham and McClung 2015; Robertson et al. 2022; Bonnot et al. 2023; Avelino et al. 2025; Ricono et al. 2025). Genes that appear strongly cold responsive at one time of day may be unchanged or oppositely regulated at another, and shifts in phase may be as biologically important as changes in overall expression magnitude. Thus, time-resolved sampling captures a major dimension of regulatory variation that would otherwise be missed.

The phase change group (PCG) coupled with the GO analysis further showed that cold acclimation retimes core biological processes into distinct diel windows. Phosphorylation/kinase signaling, intracellular protein transport, vesicle-mediated transport, RNA binding/translation, photosynthesis/light harvesting, and sulfur assimilation/redox homeostasis were recurrently enriched among PCGs, but the timing of these shifts differed among accessions. Phosphorylation/kinase signaling was repeatedly associated with transitions into or within the night, including the ZT17→ZT21 transition, suggesting that cold alters the timing of signaling processes that may coordinate acclimation responses. This interpretation is supported by evidence that phosphorylation is tightly linked to circadian regulation (Kusakina and Dodd 2012) and that plant phosphoproteomes show time-of-day variation (Uhrig et al. 2019), providing a plausible regulatory mechanism through which cold acclimation could retime specific cellular processes across the diel cycle. Photosynthesis and light harvesting genes were frequently shifted toward later daytime phases, consistent with altered regulation of light capture, photosystem function, or repair under cold (Stitt and Hurry 2002; Yarkhunova et al. 2018). In contrast, sulfur assimilation and redox-associated genes often showed similar dawn-phased expression under control conditions but diverged in cold-phase timing, suggesting that accessions may differ in how they temporally coordinate redox balance and sulfur metabolism during acclimation. Sulfur containing metabolites including cysteine and glutathione are critical for maintaining redox balance to properly respond to stress (Hossain et al. 2017; Wang et al. 2026), and while glutathione and other sulfur containing compounds do have diel cycles (Huseby et al. 2013; Cano-Yelo et al. 2026), the importance of this timing during abiotic stress remains poorly understood. Thus, the PCG/GO themes reveal both recurrent process-level diel responses and accession-specific temporal reorganization.

Circadian variation likely contributes to the diel transcriptome shifts observed here. Previous work identified that up to ∼70% of expressed genes in *B. rapa* exhibit circadian rhythmic expression (Greenham et al. 2020). Consistent with this possibility, our leaf movement assays showed that accessions differed in circadian period under control conditions and in their ability to maintain the same period under cold. Some accessions were relatively well temperature compensated, whereas others showed shortened or lengthened periods under cold, changes that could influence the apparent phase of many genes. However, because the RNA-seq experiment was conducted under diel light/dark and thermocycle conditions, we cannot fully separate direct cold responses from changes in circadian phase, temperature compensation, or diel environmental responses. Thus, some genes classified as cold responsive may reflect cold-induced reprogramming of circadian timing rather than direct transcriptional responses to cold. This distinction is important. In Arabidopsis *REVEILLE 8* mutants, which exhibit a four hour phase delay relative to wild type, accounting for the phase shift reduced the number of differentially expressed genes to only 13 (Hsu and Harmer 2012). Rather than viewing this as a limitation, these data highlight an important biological point that in field-relevant cold acclimation, stress signaling and circadian/diel regulation are intertwined. Future experiments under constant conditions, combined with acute cold treatments and clock perturbation, will be needed to distinguish direct cold-responsive genes from genes whose cold-associated expression changes arise through altered circadian timing.

A second major finding is that extensive regulatory divergence at the *B. rapa* gene level masks a more conserved regulatory architecture at the ortholog level. Pairwise comparisons of accession-level gene regulatory networks (GRNs) showed low similarity among predicted TF–target edges, and cold acclimation further reduced shared connectivity, consistent with extensive accession-specific regulatory rewiring. However, when retained *B. rapa* paralogs were collapsed to their corresponding Arabidopsis orthologs, network similarity increased substantially across accessions. This result indicates that at least part of the apparent divergence in regulatory networks reflects paralog-level diversification rather than complete loss of conserved regulatory pathways. In other words, accessions may retain related regulatory programs, but implement them through different retained paralogs. This interpretation is consistent with work in other crop species (Benoit et al. 2025) where stress responsiveness varies across paralogs (Van de Peer et al. 2017, 2021; Greenham et al. 2020; Song et al. 2020; Edger et al. 2026), and it extends that concept to inferred regulatory network structure during cold acclimation.

Integrating shared cold GRNs with PCG/GO themes connected this ortholog-level conservation to specific diel-retimed biological processes. The TF-to-theme network showed that phosphorylation/kinase signaling and RNA binding/translation/ribosome-related processes were associated with large numbers of candidate TFs, whereas photosynthesis/light harvesting, intracellular transport, vesicle trafficking, and sulfur/redox formed smaller modules. Target-level subnetworks suggest that tolerant accessions may retain specific inferred TF–target relationships within broadly shared biological processes, rather than relying on completely distinct cold-response pathways. In particular, tolerant-associated edges linked candidate regulators to a nighttime-retimed phosphorylation module and to light-harvesting/photosystem targets that shift toward later daytime expression under cold. These networks do not establish direct regulation, but they identify candidate TFs and target modules for testing how altered timing of signaling and photosynthetic regulation contributes to acclimation and freeze tolerance.

The pangenome provided the opportunity to connect these systems-level patterns to candidate cis-regulatory mechanisms through conserved noncoding sequence (CNS) analysis. CNSs can identify candidate cis-regulatory elements under evolutionary constraint (Arsovski et al. 2015; Amundson et al. 2026), and in a duplicated genome they may help distinguish which retained paralog preserves ancestral regulatory logic. Our identification of PCG-associated paralogs with CNSs shared with Arabidopsis, but absent from other members of the same *B. rapa* paralog set, suggests that differential retention or loss of ancestral cis-regulatory elements may contribute to paralog-specific diel retiming. Notable, the largest group of these CNS-linked PCG genes occurred in the ZT17->ZT21 transition, the same temporal window highlighted by the phosphorylation/kinase GO theme and tolerant-associated GRN subnetwork. This convergence among PCG timing, GO enrichment, shared GRNs, and CNS conservation provides a prioritized set of candidate genes and regulatory elements for testing how cis-regulatory variation contributes to altered timing of cold-responsive pathways.

Together, our results support a model in which cold acclimation in *B. rapa* is shaped by diel retiming of core biological processes, paralog-specific implementation of conserved regulatory programs, and accession-specific rewiring of TF-target regulatory relationships. These findings have implications for crop improvement. If tolerance depends not only on expression magnitude but also on the timing and paralog-specific regulation of stress pathways, then breeding or engineering strategies need to target cis-regulatory elements that tune when genes are expressed. The candidate TFs, PCG-associated targets, and CNS-linked paralogs identified here provide a starting point for testing whether altered temporal regulation can improve cold acclimation without broadly disrupting growth and metabolism.

## Supporting information

Supplemental Data Set 1

Supplemental Data Set 2

Supplemental Data Set 3

Supplemental Data Set 4-7

## Acknowledgements

We would like to acknowledge the loss of Ping Lou, who passed away during the analysis and preparation of this manuscript. Ping Lou assisted with the design and sampling of the RNA-seq time course experiment. The work (proposal: 10.46936/10.25585/60001202) conducted by the U.S. Department of Energy Joint Genome Institute (https://ror.org/04xm1d337), a DOE Office of Science User Facility, is supported by the Office of Science of the U.S. Department of Energy operated under Contract No. DE-AC02-05CH11231. JTL is supported by the Center for Bioenergy Innovation, which is supported by the U.S. Department of Energy, Office of Science, Biological and Environmental Research under Contract Number ERKP886.

## Author Contributions

AR, DS, AM, and KG designed the experiments. KG performed the RNA-seq time course sampling. DS, and AM, performed the freeze stress experiments. AH and AW performed the leaf movement experiment. TB, JJ, CP, JW, LB, SS, KB, JG, JS, and JTL performed the pangenome assemblies, annotations, and comparative analyses. AR, ZAM, DS, CN, and KG performed the RNA-seq and GRN analysis. YQ and ZAM performed the CNS analysis. AR, ZAM, and KG interpreted the results and wrote the manuscript.

## Data Availability

RNA-seq data is available through the NCBI GEO database. Reference genomes are available on Phytozome (https://phytozome-next.jgi.doe.gov/). All code used in this study can be found on our Github (https://github.com/greenhamlab/B.rapa-diel-cold-regulation/).

## Declaration of generative AI and AI-assisted technologies in the manuscript preparation process

During the preparation of this work, the authors leveraged Codex (5.3-mini and 5.5) and ChatGPT for code review related to data processing and to identify areas in the manuscript that could benefit from clearer language. The authors reviewed and edited the output as needed and take full responsibility for the content of the published article.

## Supplemental Figures

**Figure S1.**
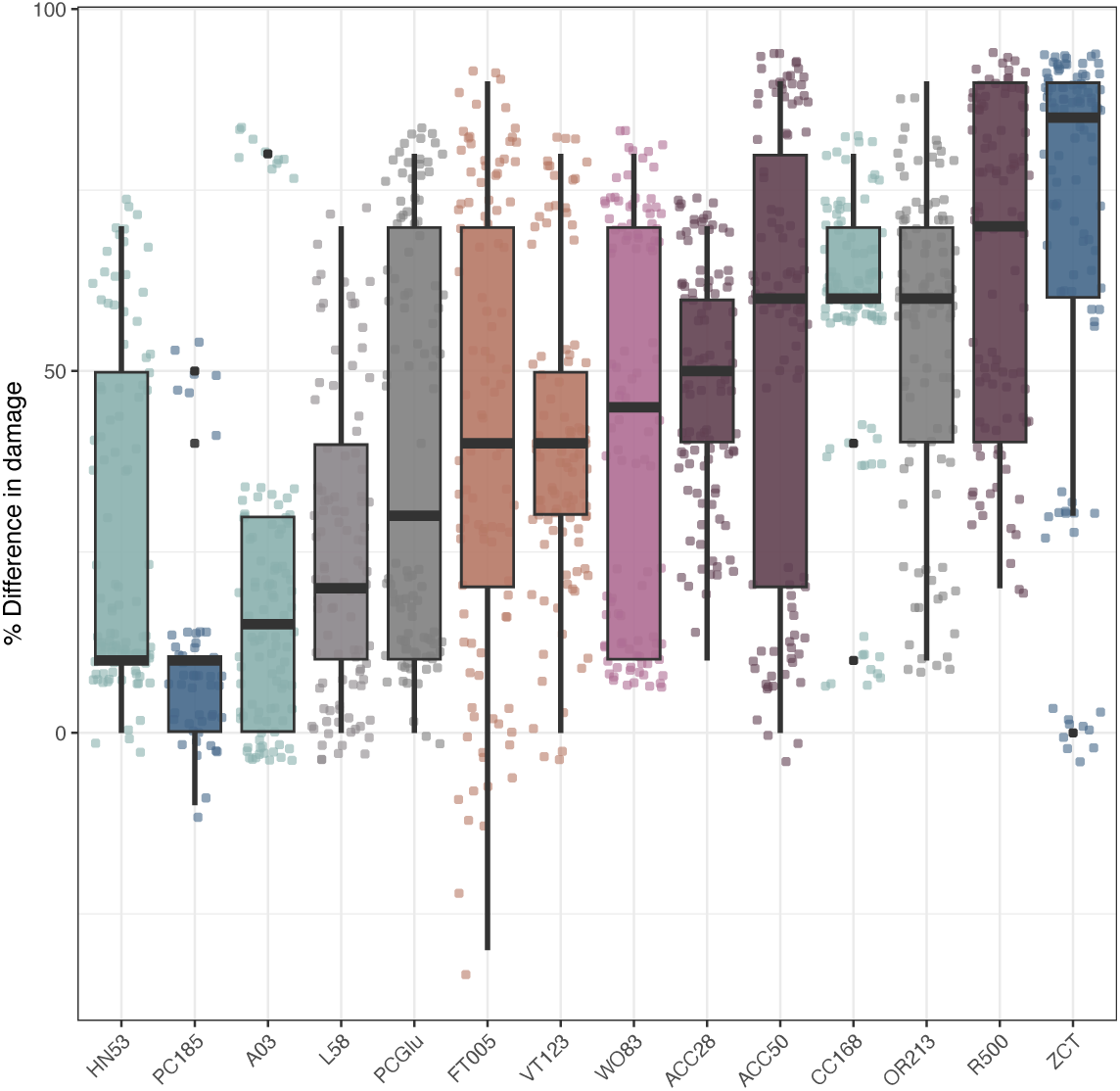
Quantitative damage score across *B. rapa* accessions under freezing stress. Data points represent individual percentages of cold responses relative to controls and are colored by morphotype; boxplots are also colored by morphotype and are ordered by mean difference in damage score from least (left) to right. Accession IDs are listed along the x-axis.

**Figure S2.**
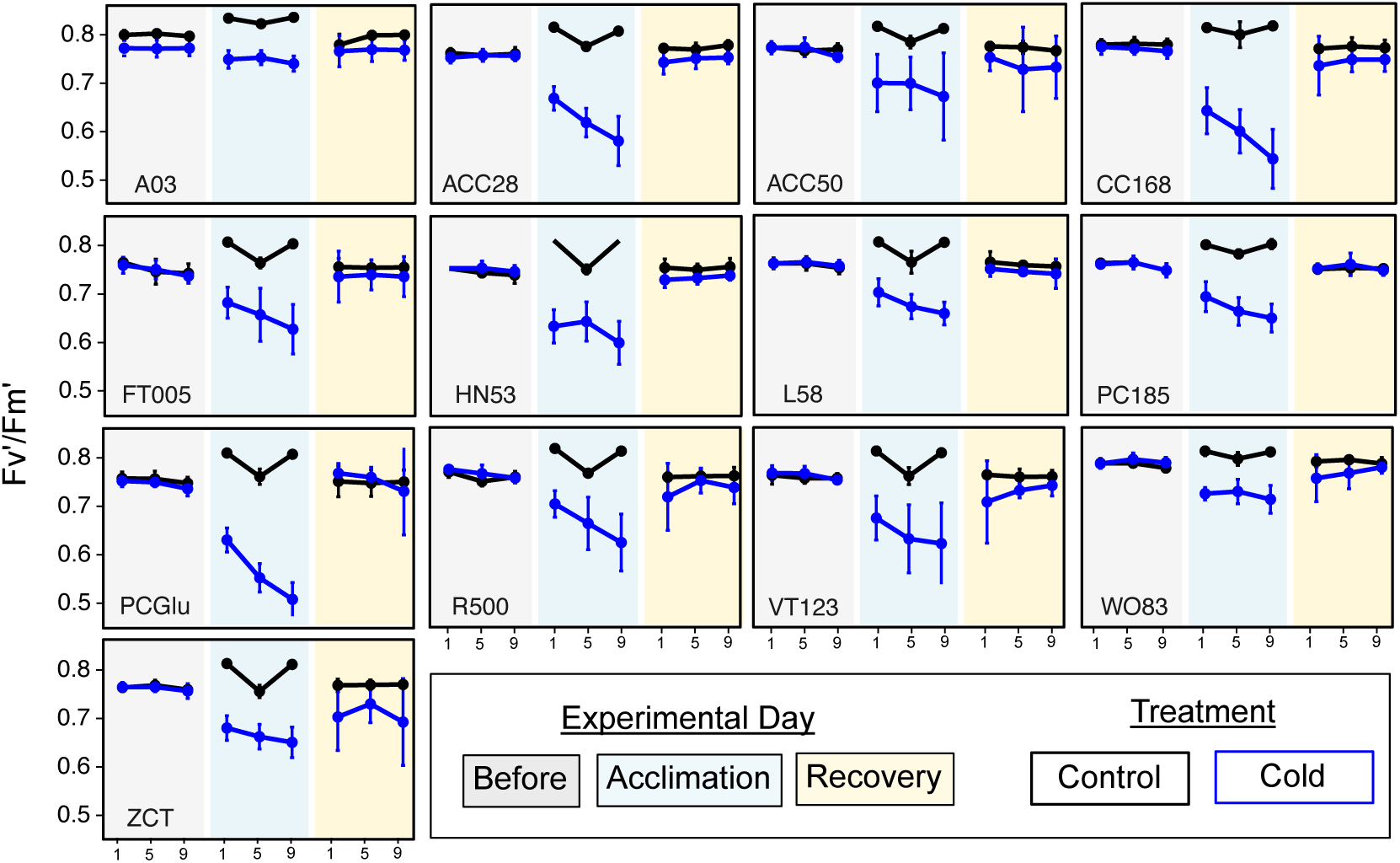
Photosynthetic efficiency among *B. rapa* accessions under freezing stress. Light adapted Photosystem II efficiency (Fv’/Fm’) was measured for each accession and treatment at ZT1, ZT5, and ZT9 before (grey background) and during (blue background) cold acclimation, and again on the recovery day following freezing stress (yellow background). accession IDs are provided on the bottom left of each panel. ZTs (Zeightgeiber; hours after lights on) are along the x-axis for each day. Lines represent the mean Fv’/Fm’ value for each group (control = black; cold = blue) while vertical bars represent the standard deviation.

**Figure S3.**
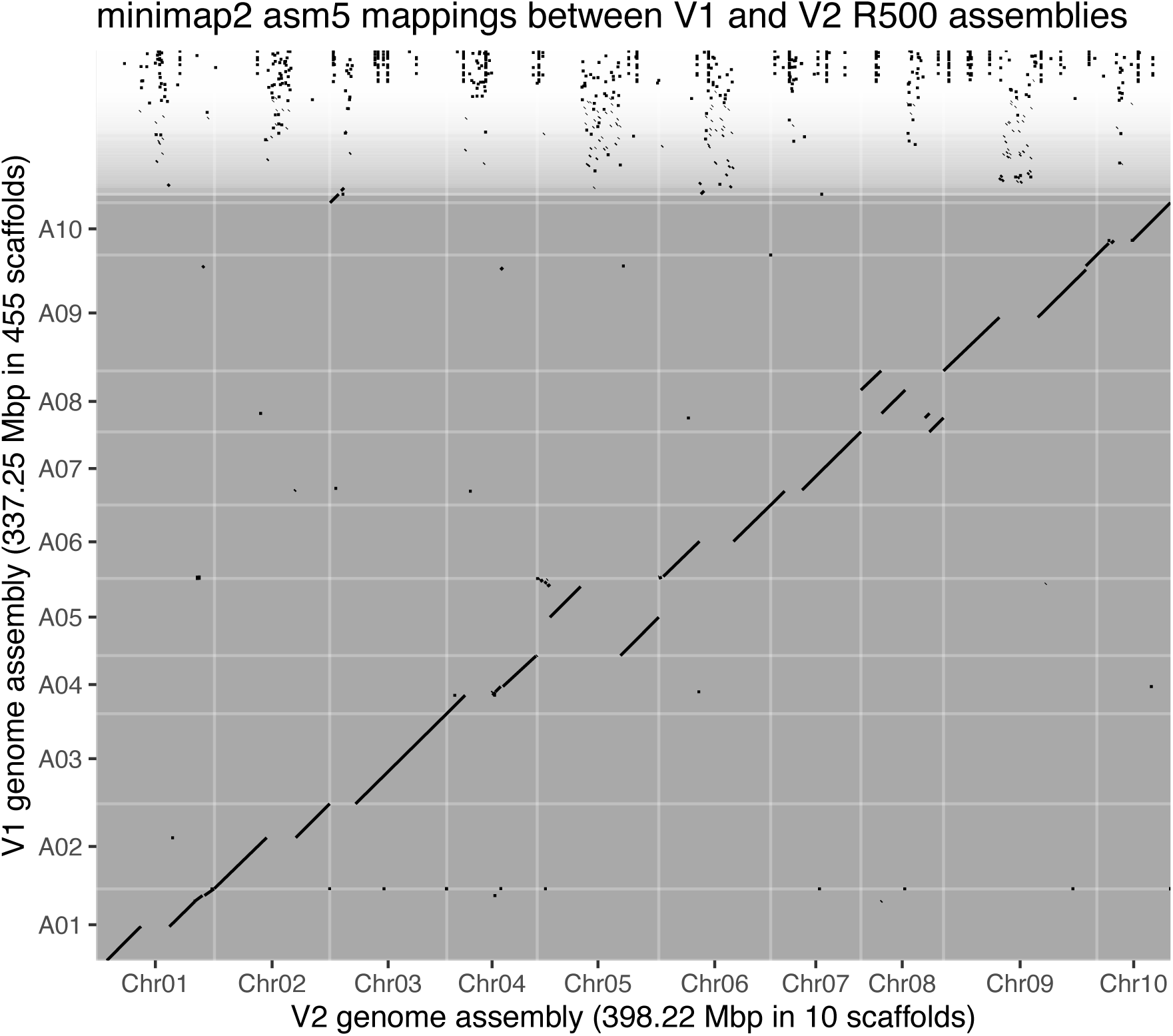
Syntenic mappings between R500 v1 and v2 genome assemblies. Minimap2 was run on settings optimized for closely related who genome alignments (-xasm5). The resulting .paf mapping file was visualized so that blocks <100kb are points and larger blocks are line segments. Only chromosome scaffolds are labeled, so all white or white-bordered facets above A10 in V1 are the unscaffolded contigs in that genome.

**Figure S4.**
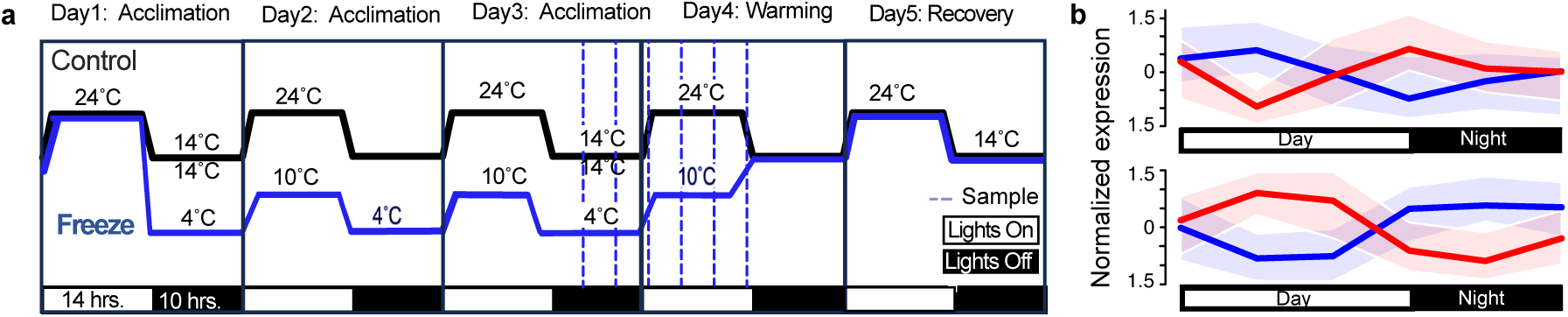
Experimental design for RNA-sequencing time course. (A) A five day time course was broken down into three cold acclimation days followed by a warming period and recovery period. Plant tissue sampling occurred at indicated ZTs(Zeitgeiber; hours since lights on) as indicated by dotted lines. Cold acclimated plant treatments are indicated by the blue line and controls by the black line. Light/dark photoperiods are indicated by white and black bars (respectively) under the plot. (B) Two representative gene expression plots for significant cold responsive genes with an advance (top) or delay (bottom) in phase following cold acclimation.

**Figure S5.**
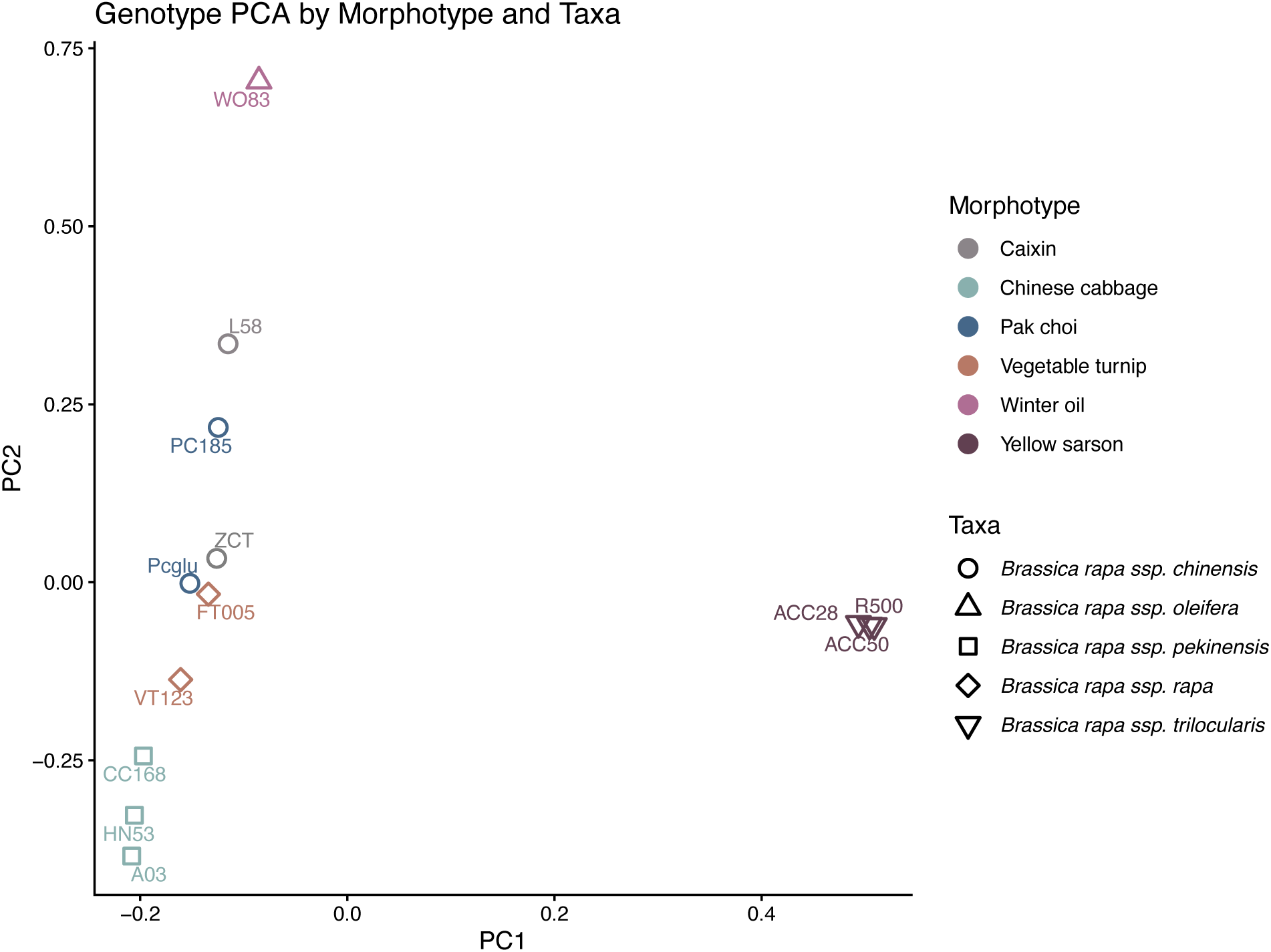
SNP-based Principal Component Analysis shows accessions cluster by morphotype. Shapes represent individual accessions as indicated next to each shape. accessions are colored by their respective morphotype.

## Supplemental Tables

**Supplemental Data Set 1. Genome Assemblies.**

**Supplemental Data Set 2. Mapping and DiPALM summary tables.**

**Supplemental Data Set 3. GO enrichment results and curated GO-theme assignments for cold-retimed phase-change genes.**

**Supplemental Data Set 4. Previously identified CNSs. Supplemental Data Set 5. CNS associations with *B. rapa* accessions.**

**Supplemental Data Set 6. Assignments of paralogs that are uniquely cold responsive in a PCG and also share a CNS with Arabidopsis.**

**Supplemental Data Set 7. Ortholog assignments.**

